# Lack of specificity in *Geobacter* periplasmic electron transfer

**DOI:** 10.1101/2022.08.12.503762

**Authors:** Sol Choi, Chi Ho Chan, Daniel R. Bond

## Abstract

Reduction of extracellular acceptors requires electron transfer across the periplasm. In *Geobacter sulfurreducens,* three separate cytoplasmic membrane cytochromes are utilized for menaquinone oxidation depending on redox potential, and at least five cytochrome conduits span the outer membrane. Because *G. sulfurreducens* produces 5 structurally similar triheme periplasmic cytochromes (PpcABCDE) that differ in expression level, midpoint potential, and heme biochemistry, separate periplasmic carriers could be needed for specific redox potentials, terminal acceptors, or growth conditions. Using a panel of marker-free single, quadruple, and quintuple mutants, the role of *ppcA* and its four paralogs was examined. Three quadruple mutants containing only one paralog (PpcA, PpcB, and PpcD) reduced Fe(III) citrate and Fe(III) oxide at the same rate and extent, even though PpcB and PpcD were at much lower levels than PpcA in the periplasm. Mutants containing only PpcC and PpcE showed defects, but were nearly undetectable in the periplasm. When expressed sufficiently, PpcC and PpcE supported wild type Fe(III) reduction. PpcA and PpcE from *G. metallireducens* similarly restored metal respiration in *G. sulfurreducens.* PgcA, an unrelated extracellular triheme *c*-type cytochrome, also participated in periplasmic electron transfer. While triheme cytochromes were important for metal reduction, sextuple Δ*ppcABCDE* Δ*pgcA* mutants still grew near wild type rates and displayed normal cyclic voltammetry profiles when using anodes as electron acceptors. These results reveal broad promiscuity in the periplasmic electron transfer network of metal-reducing *Geobacter*, and suggests an as-yet undiscovered periplasmic mechanism supports electron transfer to electrodes.

**Importance:** Many inner and outer membrane redox proteins used by *Geobacter* for electron transfer to extracellular acceptors are known to have specific functions. However, how these are connected by periplasmic redox carriers remains poorly understood. Since *Geobacter sulfurreducens* contains multiple paralogous triheme periplasmic cytochromes, each with their own unique biochemical properties and expression profiles, it has been hypothesized that each cytochrome is involved in different respiratory pathways depending on redox potential or energy conservation needs. Here we show that instead of being specific for single conditions, the many periplasmic cytochromes of *Geobacter* show evidence of being highly promiscuous. Surprisingly, while any one of 6 triheme cytochromes could support similar growth with soluble or insoluble metals, none of these were required when cells utilized electrodes. These findings could simplify construction of synthetic electron transfer pathways.

## Introduction

In anaerobic respirations where the terminal electron acceptor is too large to cross the membrane, metabolically generated electrons are routed out of the cell via a process called extracellular electron transfer (1–3). To achieve this, bacteria combine cytoplasmic membrane quinone oxidoreductases, periplasmic carriers, trans-outer membrane conduits, and conductive extracellular wires in an electrical network linking intracellular biological reactions to extracellular events (2, 4). These unique electron transport chains alter metal redox states, directly exchange electrons with other bacteria, and provide new tools for bioelectronic applications (5–8).

*Geobacter sulfurreducens* is a model of extracellular electron transfer due to its ability to reduce acceptors such as Fe(III), Mn(IV), U(VI), V(V), Tc(VII), and electrodes (9). A central question raised by this versatility involves whether different redox proteins are required for each acceptor. To date, it is known that three separate inner membrane quinone oxidases are utilized depending on the redox potential of the terminal acceptor (10–13). In contrast, of five characterized outer membrane-spanning cytochrome conduits, specific complexes are linked to reduction of each type of metal or surface (14). Separate multiheme cytochrome nanowires are also used during growth with different terminal electron acceptors (15–17).

Some electron transfer mechanism is required to connect this array of inner and outer membrane proteins, as they are electrically separated by the ~30 nm-wide periplasm and cell wall (18, 19). Periplasmic redox carriers are often small promiscuous proteins able to form weak complexes and facilitate rapid electron transfer with multiple structurally unrelated proteins (20–25). A well-studied candidate for this role in *G. sulfurreducens* is the highly abundant 10 kDa triheme *c*-type cytochrome PpcA. Deletion of *ppcA* causes significant defects in Fe(III) reduction (26, 27), and purified PpcA forms functional low affinity complexes with multiple periplasmic redox partners (23, 28, 29). However, *G. sulfurreducens* also produces four paralogs (PpcB, PpcC, PpcD, and PpcE) with highly similar heme arrangements and protein folds, and genetic evidence indicates these proteins can influence electron transfer (30, 31). For example, in growth-independent assays, deletion of *ppcA* impacts, but does not eliminate, insoluble Fe(III) oxide reduction, and has an even smaller effect on soluble Fe(III) citrate reduction (26). While PpcB can be more abundant during Fe(III) citrate growth, deletion of *ppcB* increases Fe(III) citrate reduction rates (31, 32). PpcE is rarely detected, but deletion of *ppcE* is reported to increase Fe(III) reduction rates (32).

Biochemical differences further suggest unique roles for Ppc paralogs. PpcA has two higher potential hemes near −100 mV vs. Standard Hydrogen Electrode (SHE), and only one below −150 mV, while other paralogs show the opposite pattern (33). PpcC exists in two unique conformations, depending on its oxidation state (34). PpcA and PpcD display a phenomenon coined the ‘Redox-Bohr’ effect, where reduction shifts the pKa of a heme propionate side chain more than one pH unit (35, 36). When certain PpcA and PpcD hemes are reduced, this shift can cause uptake of a proton. Protonation makes heme oxidation less favorable by ~ 50 mV, but if an adequate acceptor is available, oxidation drives proton release. At a cost of about 50 mV/e^-^, this could transport a proton across the periplasm, in a cycle proposed to aid energy generation when cells use PpcA or PpcD (35, 36).

While it is known that expression of *ppcA* alone in a mutant lacking all 5 paralogs will fully restore wild type Fe(III) citrate reduction, data supporting functional roles for the remaining periplasmic carriers is incomplete and sometimes contradictory (27). In this work, a panel of marker-free single, quadruple, and quintuple *ppc* deletion mutants were constructed. Some quadruple mutants containing only one paralog showed defects in Fe(III) reduction, but these changes were correlated with low periplasmic cytochrome abundance. Standardizing periplasmic levels using the *ppcA* promoter, ribosomal binding site (RBS), and/or signal peptide showed for the first time that any *ppc* cytochrome, when expressed sufficiently, will support wild type reduction of both Fe(III) citrate and Fe(III) oxide. PpcA and PpcE from *G. metallireducens* were also capable of restoring metal respiration in *G. sulfurreducens.* No significant differences in growth rate, extent of reduction, or use of specific acceptors could be found in cells using any form of PpcA or PpcD, compared to PpcB, PpcC or PpcE. PgcA, an unrelated extracellular triheme *c*-type cytochrome (37), was also shown to contribute to periplasmic electron transfer. Surprisingly, while triheme cytochromes were absolutely essential for wild type level metal reduction, the sextuple deletion mutant lacking *ppcABCDE* and *pgcA* still grew near wild type rates when using anodes as electron acceptors. These results reveal broad promiscuity in this periplasmic cytochrome family, and show that electron transfer to electrodes can use a Ppc-independent mechanism.

## Results

### Single deletions of Ppc-family cytochrome genes do not affect soluble or insoluble Fe(III) reduction

*G. sulfurreducens* encodes five homologous ~10 kDa triheme periplasmic cytochromes which range in pairwise identity from 46 - 77%. To test if any of these cytochromes were necessary for reduction of Fe(III) citrate or Fe(III) oxide, markerless single deletion mutants lacking *ppcA, ppcB, ppcC, ppcD,* or *ppcE* were compared to wild type (Fig 1A, 1B). None of these single deletion mutants exhibited a defect in Fe(III) reduction rate, or in the final extent of reduction for either Fe(III) form, indicating that no single gene was essential under the conditions tested.

**Fig 1.**
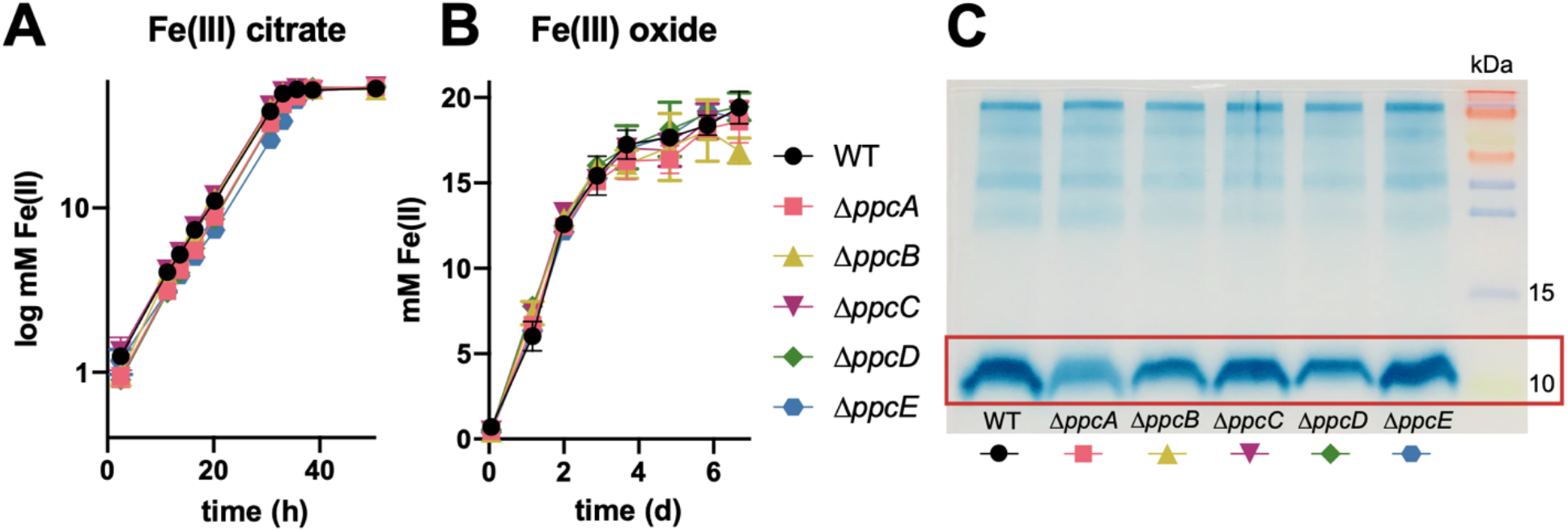
Soluble Fe(III) citrate and insoluble Fe(III) oxide reduction is not affected in *G. sulfurreducens* mutants lacking single Ppc-family cytochromes. (A) Fe(III) citrate reduction. (B) Fe(III) oxide reduction. (A) and (B) represent mean ± standard deviations with 3 technical replicates. (C) Heme-staining of periplasmic fractions from WT and single deletion mutants, representative of two independent experiments.

Prior transcriptional and proteomic studies have shown that other PpcA paralogs are always expressed and present in the periplasm (31). Consistent with this, when periplasmic proteins were obtained, 10 kDa cytochromes could still be detected in each *G. sulfurreducens* single mutant (Fig 1C). Deletion of *ppcA* caused the greatest decrease in the abundance of the pool, followed by Δ*ppcD* and Δ*ppcB.* In contrast, Δ*ppcC* or Δ*ppcE* showed little decrease in the abundance of 10 kDa periplasmic cytochromes.

### Quadruple mutants reveal PpcA, PpcB and PpcD are equally able to support WT Fe(III) reduction

Quadruple markerless deletion strains were then constructed to better isolate the roles of individual cytochromes. Mutants lacking four *ppc*-homologs were labeled according to the gene still remaining in its original genomic context. For example, Δ*ppcBCDE* was referred to as *ppcA*^+^, Δ*ppcABCD* as *ppcE*^+^, and the strain lacking all five (Δ*ppcABCDE*) was referred to as Δ*ppc*5.

Three quadruple deletion strains (*ppcA*^+^, *ppcB*^+^, and *ppcD*^+^) retained wild type rates and extents of both Fe(III) citrate and Fe(III) oxide reduction. In contrast, *ppcC*^+^ and *ppcE*^+^ showed strong defects with both acceptors. Deletion of all five *ppc* cytochromes further diminished, but did not eliminate, Fe(III) reduction (Fig 2A, 2B). This Δ*ppc*5 strain grew 70% slower than wild type with Fe(III) citrate (Fig 2).

**Fig 2.**
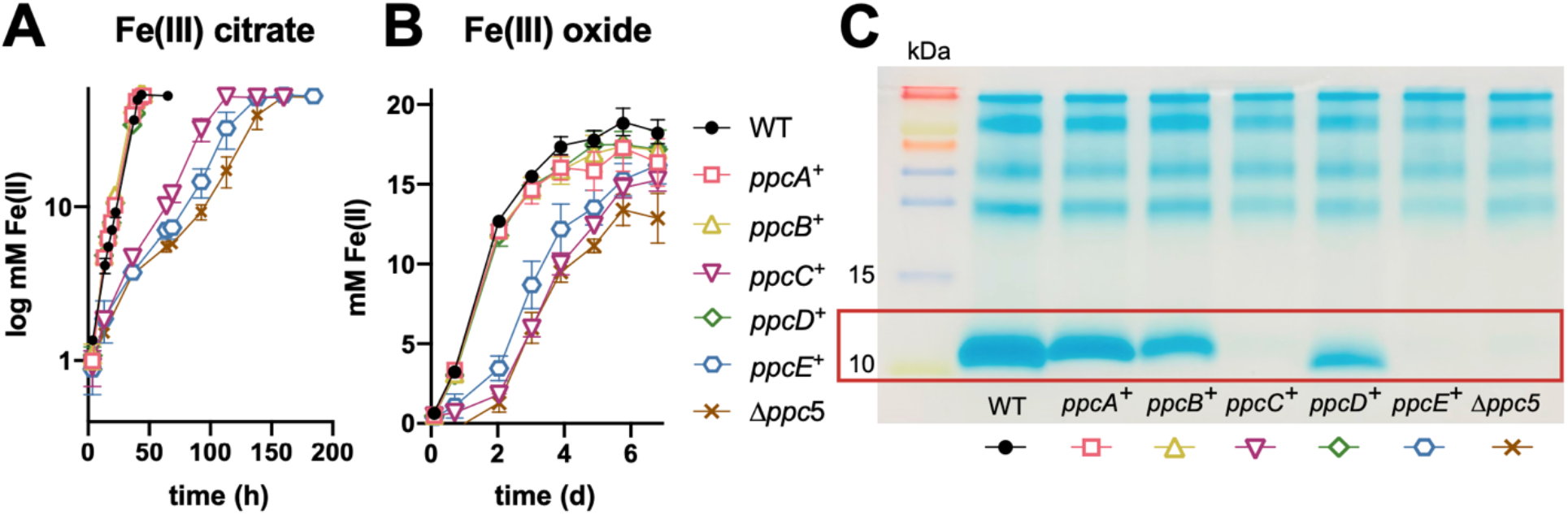
Single periplasmic cytochromes are to support Fe(III) citrate and Fe(III) oxide reduction, based on quadruple and quintuple *ppc* deletion strains. (A) Fe(III) citrate reduction (n=3 independent replicates). (B) Fe(III) oxide reduction (n=2) (C) Heme-staining of the periplasmic fraction showing nearly undetectable 10 kDa cytochrome in *ppcC*^+^ and *ppcE*^+^ mutants, representative of five independent experiments.

When periplasmic proteins from quadruple and quintuple mutants were compared (Fig 2C), 10 kDa cytochromes were only detected in *ppcA*^+^, *ppcB*^+^, and *ppcD*^+^, the strains demonstrating wild type growth. In contrast, cytochrome at 10 kDa was nearly undetectable in strains with strong defects (*ppcC*^+^, *ppcE*^+^, and Δ*ppc*5). Consistent with data from single mutants (Fig 1C), where deletion of *ppcA* caused the largest decrease in cytochrome abundance, the intensity of *ppcA*^+^ was most intense among the quadruple deletion strains, followed by *ppcD*^+^ and *ppcB*^+^. This showed that even cytochromes with much lower native abundance, such as PpcB and PpcD, could still support metal reduction similar to the more abundant PpcA.

These data also suggested that the primary explanation for the failure of *ppcC*^+^ and *ppcE*^+^ strains to reduce Fe(III) could be related to their low abundance, rather than any biochemical specificity. As these periplasmic fractions were routinely obtained from fumarate-grown cells, periplasmic fractions were also collected from all mutants grown in Fe(III) citrate (Fig S2). Consistent with prior transcriptional studies reporting few differences in *ppc* paralog expression levels between these two conditions (Fig S3), *ppcA*^+^, *ppcB*^+^, and *ppcD*+ strains contained detectable 10 kDa cytochrome, while *ppcC*^+^, *ppcE*^+^, and Δ*ppc*5 did not.

### PgcA, previously identified as an extracellular *c*-type cytochrome, also contributes to periplasmic electron transfer

The finding that the Δ*ppc*5 strain had significant residual extracellular electron transfer ability was unexpected. We designed enrichments for spontaneous mutants upregulating or increasing use of cryptic mechanisms in strains lacking some or all Ppc-family cytochromes. While incubations of quintuple Δ*ppc*5 strains did not readily yield faster-growing suppressor strains, a slow-growing strain lacking the three most abundant Ppc-cytochromes (Δ*ppcABD*), evolved accelerated Fe(III) citrate reduction. Re-isolation and re-sequencing identified a single 9 bp in-frame deletion within a gene encoding *pgcA,* a triheme cytochrome unrelated to any *ppc* homologs (Fig 3A).

**Fig 3.**
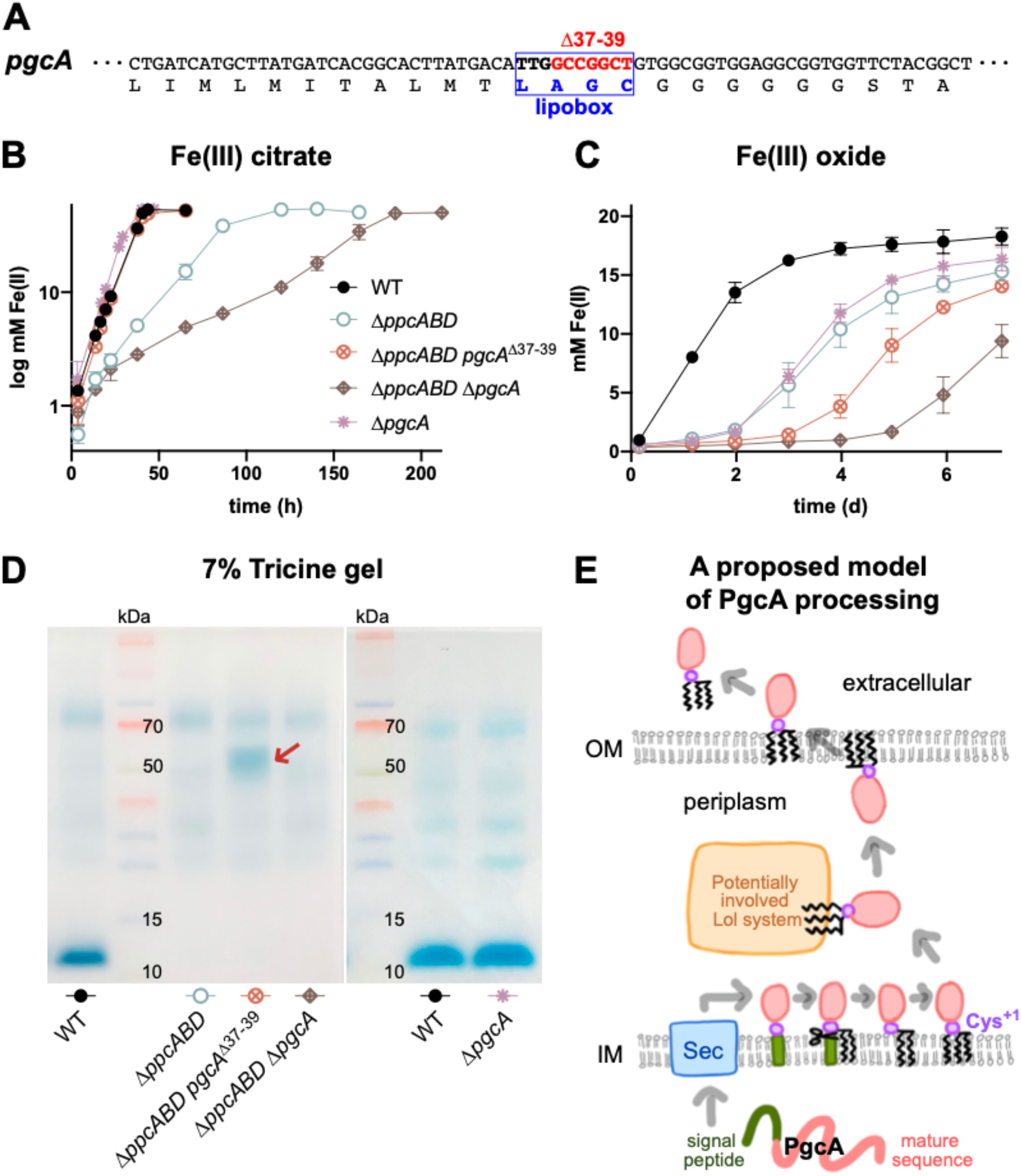
Evidence that PgcA participates in periplasmic electron transfer. (A) Partial amino acid sequence of PgcA containing the lipobox domain (blue) and deleted nucleotides (red) in Δ*ppcABD pgcA*^Δ37-39^. Fe(III) reduction (B,C) and periplasmic cytochrome abundance (D) of the Δ*ppcABD pgcA*^Δ37-39^ strain, showing increased Fe(III) citrate reduction by cells retaining PgcA in the periplasm, and expected Fe(III) oxide defect. (E) Proposed model of PgcA processing, from (46).

PgcA is an extracellular triheme *c*-type cytochrome with a putative lipoprotein signal sequence (37). As the suppressor strain deleted three amino acids (A-G-C) within the N-terminus ‘lipobox’ that is typically the target of acylation before cleavage by the lipoprotein-specific signal peptidase (Lsp/SpII) (45), this strain was designated Δ*ppcABD pgcA*^Δ37-39^. We hypothesized that the mutated signal peptide was inhibiting translocation of PgcA out of the periplasm (Fig 3E).

Periplasmic proteins from the suppressor strain Δ*ppcABD pgcA*^Δ37-39^ contained a new abundant cytochrome near the molecular weight of PgcA (50 kDa) (Fig 3D and Fig S4). Ultracentrifugation of periplasmic fractions did not cause any significant change in the abundance of cytochromes, confirming that membrane-bound cytochromes were not present in these periplasmic fractions (Fig S1). Deletion of the mutant *pgcA*^Δ37-39^ from Δ*ppcABD pgcA*^Δ37-39^ eliminated this 50 kDa periplasmic cytochrome (Fig 3D and Fig S4), and produced a strain even more impaired than the parent Δ*ppcABD* (Fig 3B).

When native *pgcA* was deleted from the Δ*ppc*5 background, the resulting Δ*ppc*5 Δ*pgcA* demonstrated the lowest rate of metal reduction, less than 10% of wild type with Fe(III) citrate (Fig 4). These lines of evidence support a model where a change in PgcA localization increased its periplasmic abundance to rescue wild type periplasmic electron transfer.

**Fig 4.**
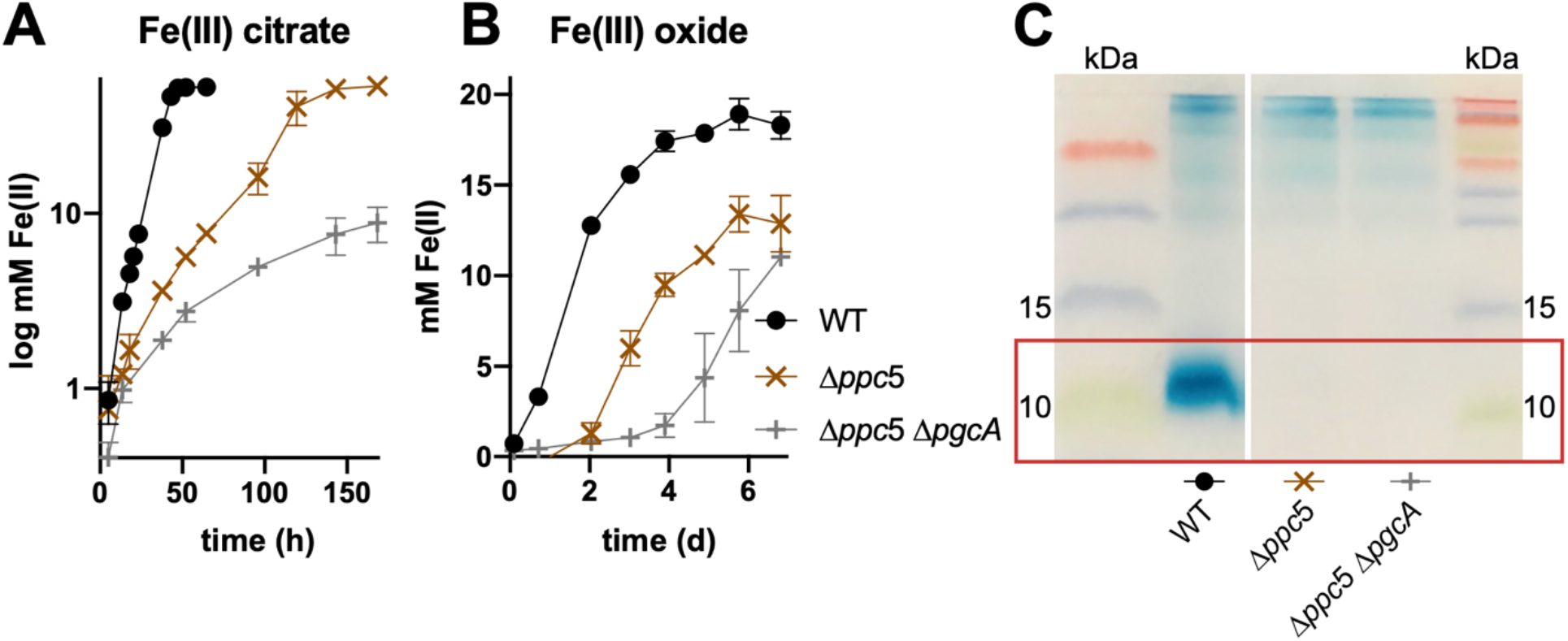
Decrease in residual Δ*ppc*5 electron transfer ability by deletion of *pgcA.* (A) Fe(III) citrate reduction. (B) Fe(III) oxide reduction. Curves in (A) and (B) are representative of three independent biological replicates. (C) Heme-staining of the periplasmic fraction (16% tricine), representative of two independent experiments.

While PgcA is not essential for Fe(III) citrate reduction in wild type cells, extracellular PgcA is critical for rapid insoluble metal oxide reduction (37). According to the hypotheses that PgcA is retained in the periplasm of the Δ*ppcABD pgcA*^Δ37-39^ strain, cells should have the same defect as Δ*pgcA* when Fe(III) oxide is the acceptor. Consistent with this model, both Δ*ppcABD pgcA*^Δ37-39^ and Δ*ppc*5 Δ*pgcA* strains reduced Fe(III) oxide poorly (Fig 3B and 3C). Together, these data suggest that, as PgcA is being processed in the periplasm of wild type cells, it can participate in periplasmic electron transfer. As *pgcA* is typically more highly expressed during reduction of insoluble metal oxides (31), it likely only plays a minor role during reduction of soluble compounds.

### Engineering *ppcC* and *ppcE* for increased periplasmic abundance shows these cytochromes can also support WT Fe(III) reduction

Strains containing only *ppcC* or *ppcE* under control of their native promoters reduced Fe(III) poorly and demonstrated a lack of 10 kDa periplasmic cytochromes in periplasmic extracts (Fig 2, Fig S3). Thus, we sought to more fairly test the properties of individual cytochromes by developing a standardized expression method. Beginning with the Δ*ppc5 ΔpgcA* mutant containing the lowest background levels of metal reduction (Fig 4), *ppcC* and *ppcE* were cloned downstream of the *ppcA* promoter-RBS sequence, as *ppcA* is one of the most highly expressed genes in *G. sulfurreducens* (13). As controls, *ppcA* and *ppcB* were cloned downstream of the same promoters using the identical protocol. All constructs were integrated in a neutral site downstream of *glmS* (Fig 5A).

**Fig 5.**
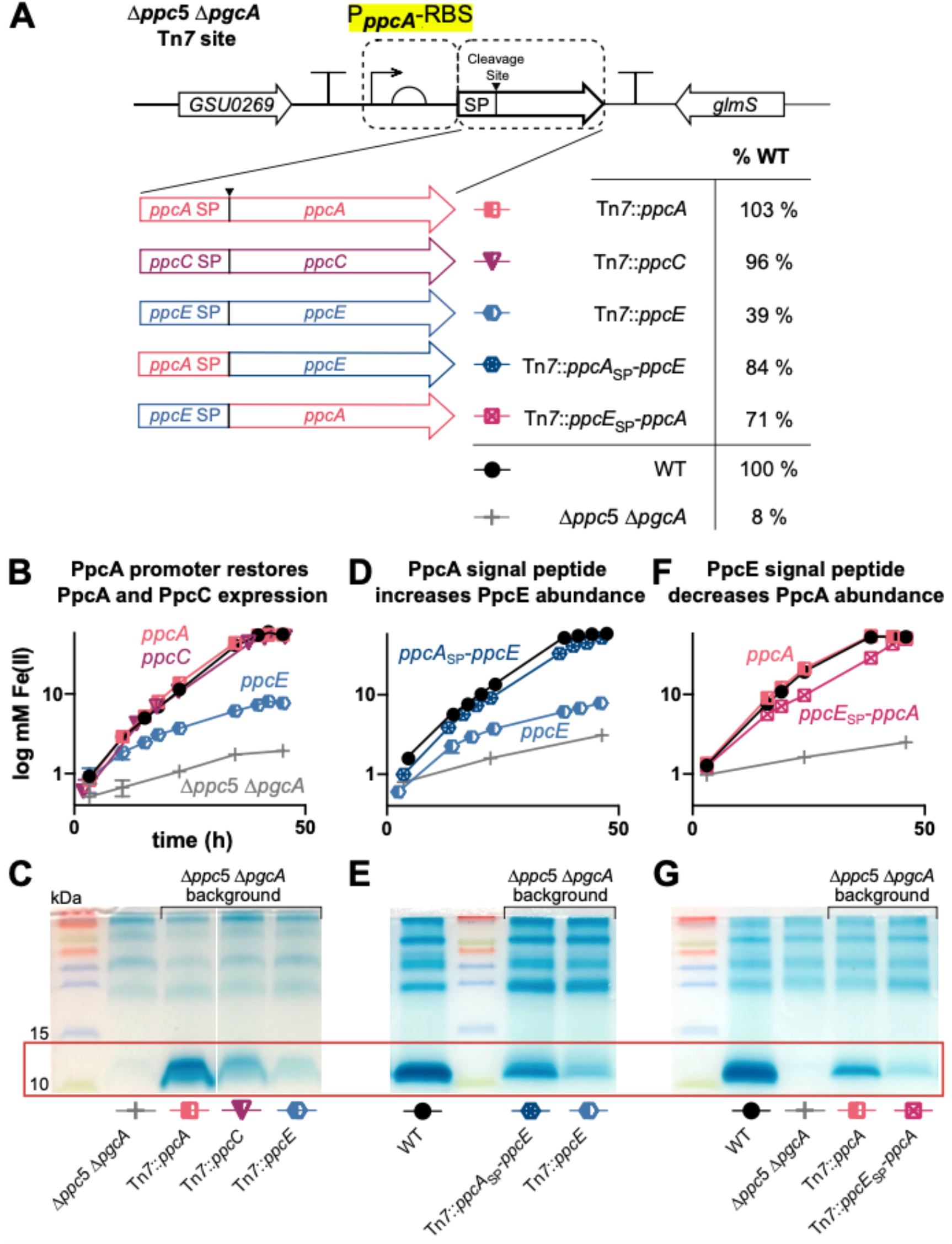
Fe(III) reduction by PpcC and PpcE-containing strains is similar when periplasmic cytochrome abundance is restored. (A) Cytochrome re-expression strategy, combining the *ppcA* promoter and RBS (highlighted) with different signal peptides (SP). (B,D,F) Fe(III) citrate reduction (n=3) (C,E,G) Heme-stained periplasmic fractions. While PpcC abundance increased with use of the *ppcA* promoter (B,C), PpcE required both the *ppcA* promoter and PpcA signal peptide (D, E). Fusion of the PpcE signal peptide to PpcA decreased PpcA abundance (F, G).

Expression of *ppcA* in this system successfully rescued wild type Fe(III) reduction and produced abundant periplasmic PpcA in Δ*ppc*5 Δ*pgcA* (Fig 5B, 5C). Similar results were obtained when *ppcB* was expressed using the same strategy (Fig S5). When *ppcC* was expressed under control of this *ppcA* promoter system, wild type Fe(III) reduction was also rescued, and moderate levels of periplasmic PpcC were detected (Fig 5B, 5C). However, expression of *ppcE* under the same conditions (Δ*ppc*5 Δ*pgcA Tn*7::*ppcE*) only partially improved metal reduction, and produced barely detectable periplasmic cytochrome around 10 kDa (Fig 5D, 5E).

Multiple modifications were tested to identify the cause of low PpcE abundance. Recoding *ppcE* to eliminate rare *G. sulfurreducens* codons did not improve the abundance of PpcE or rescue wild type Fe(III) citrate reduction (Fig S6). However, when the PpcE signal peptide was replaced with the PpcA signal peptide sequence (*ppcA*sp-*ppcE*), metal reduction improved to near wild type, and PpcE in the periplasm increased (Fig 5D, 5E). To further test the hypothesis that PpcE signal peptides affected periplasmic protein levels, we generated a chimeric protein replacing the signal peptide of PpcA with that of PpcE (*ppcE*sp-*ppcA*) (Fig 5F and 5G). The PpcE signal peptide strongly decreased PpcA abundance, even though the gene remained under control of the *ppcA* promoter-RBS (Fig 5G). This decrease only caused a small defect in Fe(III) citrate reduction (Fig 5F), again showing that large amounts of a Ppc cytochrome were not needed for rapid electron transfer.

Finally, PpcC and PpcE constructs were introduced into the Δ*ppc*5 background (as PgcA is necessary for Fe(III) oxide reduction) to test their ability to participate in Fe(III) oxide reduction. Both the PpcC and PpcE re-expression strains regained Fe(III) oxide reduction to rates similar to the wild type (Fig 6A). In addition, periplasmic levels of each cytochrome in the Δ*ppc*5 background were similar to what was obtained in the Δ*ppc*5 Δ*pgcA* background (Fig 6B). These combined results show that PpcB, PpcC, PpcD, and PpcE can support wild type rates and extents of both soluble and insoluble metal reduction in *G. sulfurreducens.*

**Fig 6.**
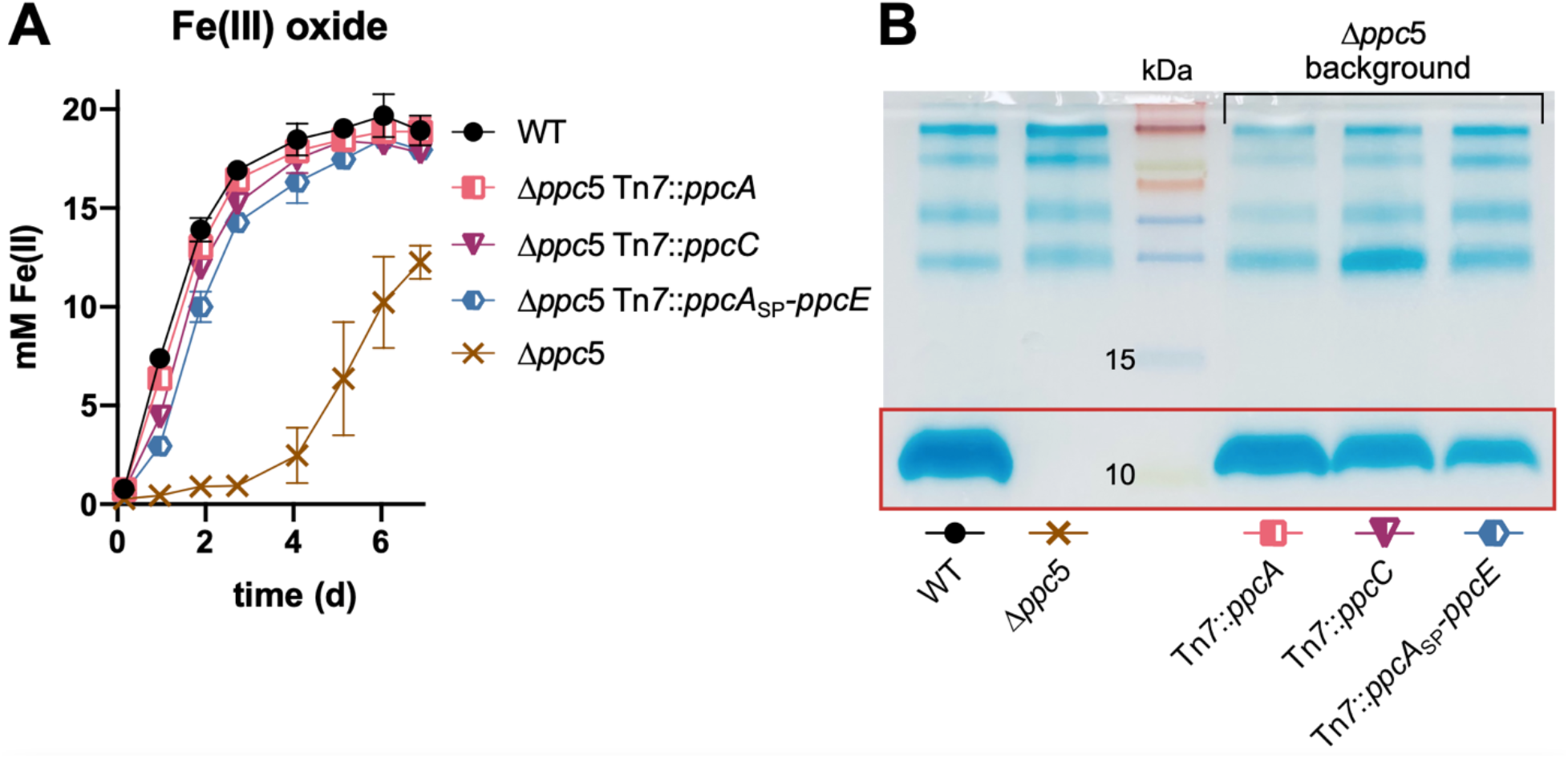
Expression of *G. sulfurreducens ppcC* and *ppcE* cytochromes in Δ*ppc*5 also rescues Fe(III) oxide reduction. (A) Fe(III) oxide reduction. (B) Heme-staining of periplasmic fraction. The gel is representative of two independent experiments.

### Expression of periplasmic cytochromes from other organisms

As all Ppc paralogs, and even an unrelated extracellular cytochrome, could support periplasmic electron transfer during metal reduction in *G. sulfurreducens,* a panel of homologs were tested for their ability to be targeted to the periplasm and restore electron transfer in the Δ*ppc*5 Δ*pgcA* background. These included the PpcA and PpcE homologs from *G. metallireducens* (81% and 62% identity to PpcA in *G. sulfurreducens),* a structurally related tetraheme *c*-type cytochrome from *Desulfovibrio vulgaris* (Fig S7C), and the tetraheme CctA (Fig S7D) involved in *Shewanella oneidensis* periplasmic electron transfer (Fig 7).

**Fig 7.**
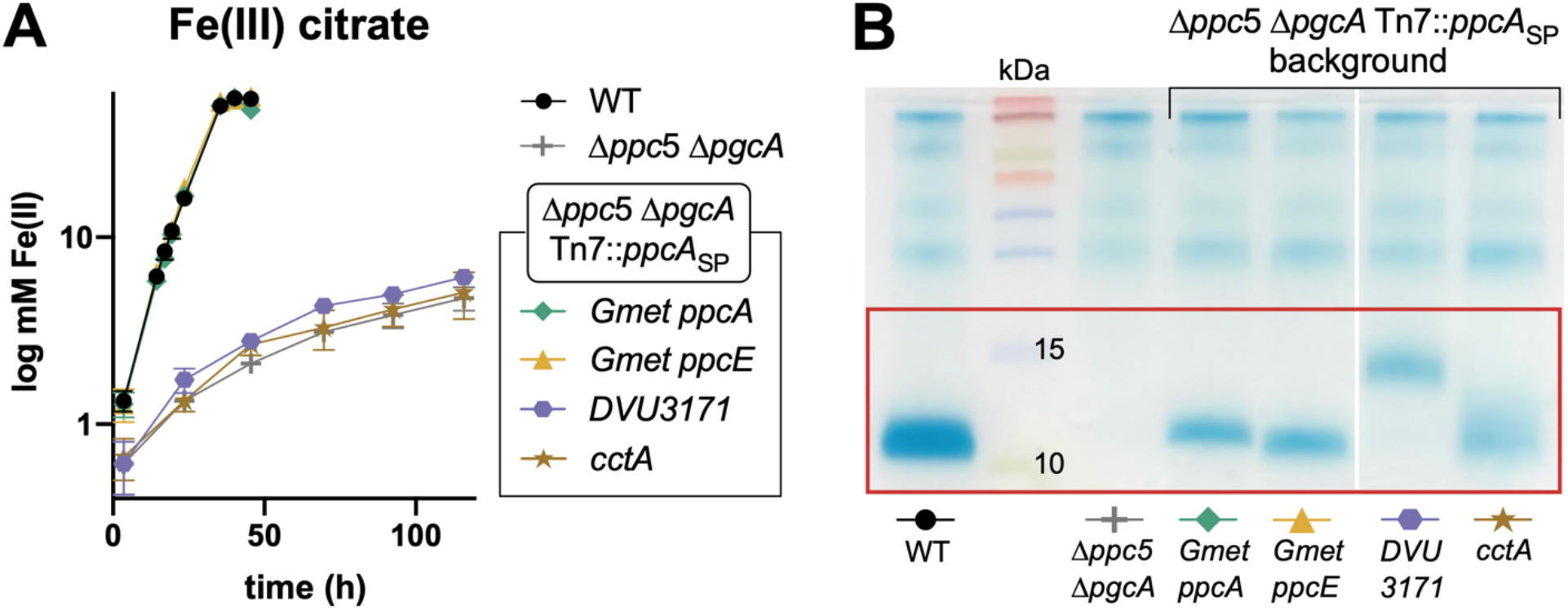
Successful heterologous expression of periplasmic cytochromes from other species in *G. sulfurreducens.* (A) Fe(III) citrate reduction. Curves in (A) are representative of two independent experiments. (B) Heme-staining of periplasmic fractions showing proper localization of introduced cytochromes.

All heterologous constructs used the *G. sulfurreducens ppcA* promoter, ribosomal binding site, and signal peptide sequence, and were detected in the periplasm (Fig 7B). However, only the homologs from *G. metallireducens* fully rescued Fe(III) citrate reduction in Δ*ppc5* Δ*pgcA* (Fig 7A). CctA and DVU3171 did not improve Fe(III) citrate reduction.

### Deletion of all six periplasmic cytochromes has little effect on electrode reduction

In every experiment up to this point, as long as the cytochrome was detectable in the periplasmic fraction, *ppcA, ppcB, ppcC, ppcD,* or *ppcE* supported comparable rates and extents of reduction, and the small amount of background electron transfer activity observed in the absence of these five genes could be attributed to *pgcA.* Based on these results, extracellular respiration by *G. sulfurreducens* requires at least one of these periplasmic cytochromes for wild type level reduction, and the Δ*ppc*5 Δ*pgcA* strain would be expected to be highly defective with all other extracellular electron acceptors.

Unexpectedly, the Δ*ppc*5 Δ*pgcA* strain grew only 8% slower when using a poised electrode (+0.24 vs. SHE) as the sole electron acceptor, and reached the same maximal current as the wild type (Fig 8A). During the exponential phase of growth, the Δ*ppc*5 Δ*pgcA* mutant actually produced current 12% faster than the wild type (expressed as μA produced per μg protein, n=4, measured at 100 μA·cm^-2^). This rate of current production coupled with a slightly slower growth rate predicted a minor defect in growth yield. Dividing the amount of biomass on electrodes at 100 μA·cm^-2^ by the integrated amount of current produced by the time of harvest was consistent with a 13% reduction in apparent yield of Δ*ppc*5 Δ*pgcA* (as protein/coulomb) (Fig 8A).

**Fig 8.**
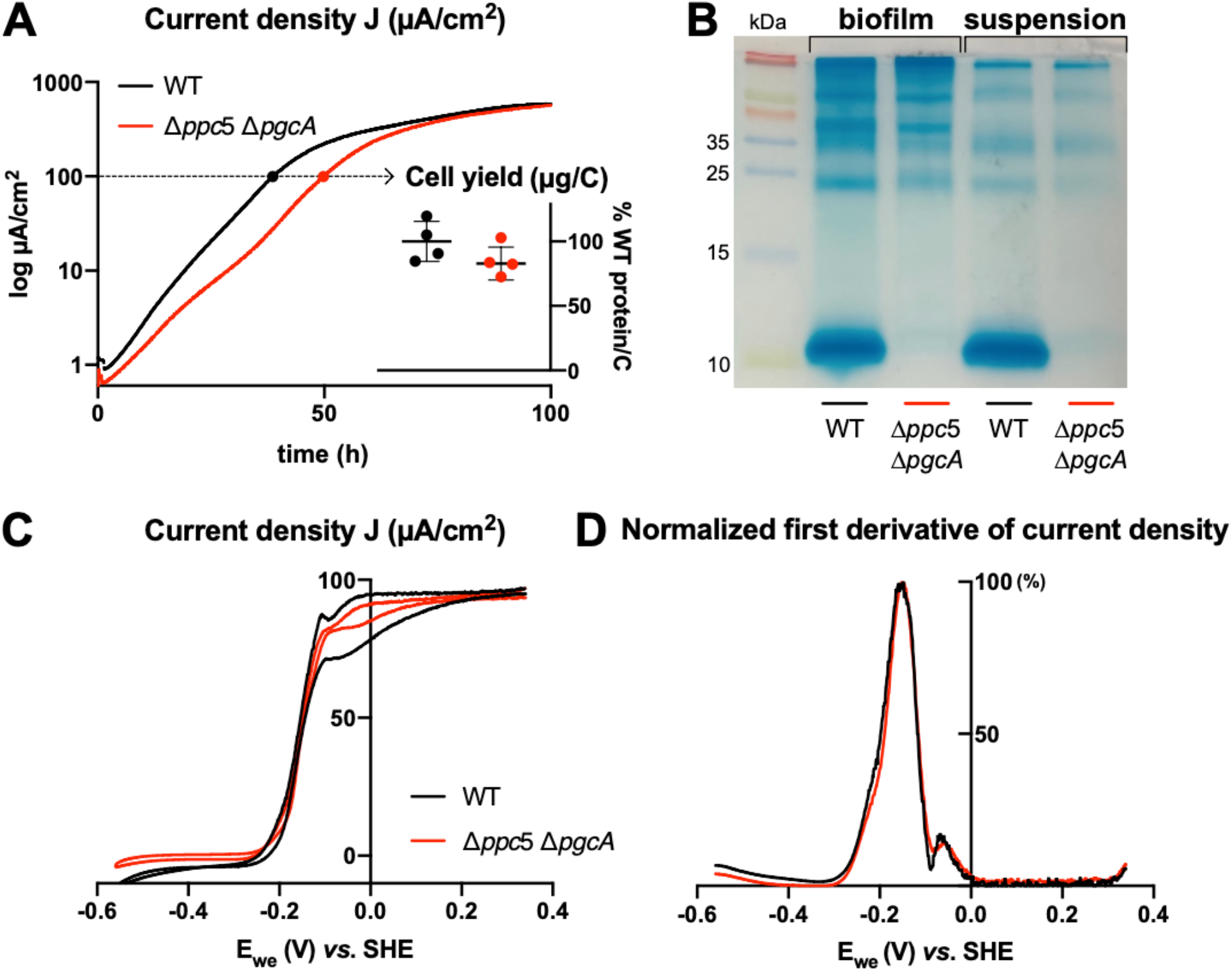
No known periplasmic cytochrome is important for electrode reduction. Comparison of wild type vs. Δ*ppc*5 Δ*pgcA* grown with an anode poised at +0.24 V vs SHE. (A) Chronoamperometry of WT and Δ*ppc*5 Δ*pgcA* (B) 16% tricine gel of osmotic shock periplasmic fractions from biofilm and planktonic cells after 4 days of electrode growth. (C) Cyclic voltammetry of wild type and Δ*ppc*5 Δ*pgcA*. (D) First derivative of data from (C).

Periplasmic fractions recovered from both planktonic and anode biofilm cells did not reveal induction of any new periplasmic cytochromes that could explain the unexpected growth of Δ*ppc*5 Δ*pgcA* with electrodes (Fig 8B). Cyclic voltammetry showed that the characteristic onset and midpoint potentials were similar in wild type and Δ*ppc*5 Δ*pgcA*, indicating electron transfer across a range of redox potentials was unaffected (Fig 8C and 8D). The only qualitative difference observed was a reduced hysteresis between forward and reverse scans, a feature that could reflect lower electron storage capacity in cells lacking the normally highly abundant Ppc family cytochromes. Otherwise, there was no evidence that any of the six triheme cytochromes removed from this strain were necessary for electron transfer to electrodes.

### Evidence for triheme cytochromes being necessary for oxidative stress protection

Previous research reported a transient interaction between PpcA and the diheme cytochrome *c* peroxidase MacA (12, 13). If periplasmic cytochromes provide reducing power to peroxidases, mutants should have increased sensitivity to H_2_O_2_ stress. In lawns of cells exposed to 3% H_2_O_2_-soaked filter discs, the zone of inhibition was unchanged for any single *ppc* deletion mutant compared to wild type cells (Fig S5). Mutants that that lacked most periplasmic cytochromes (*ppcC*^+^, *ppcE*^+^, Δ*ppc*5, and Δ*ppc*5 Δ*pgcA*) exhibited detectable larger zones of inhibition (Fig S8). These data were consistent with multiple Ppc-family cytochromes, as well as PgcA, aiding H_2_O_2_ detoxification.

## Discussion

Every *Geobacter* genome contains between 4-6 PpcA paralogs that can share similar heme packing and backbone structures, but have significant differences in redox potentials, microstates of partial oxidation, protonation behaviors, and surface charges near solvent-exposed hemes (30, 35). PpcA homologs from *G. sulfurreducens* and *G. metallireducens* show such large differences in midpoint potential and heme oxidation order that the two cytochromes are proposed to interact with different redox partners. In this study, we could find no direct evidence that these biochemically different proteins had unique roles, redox potentials, partners, or energetic benefits during reduction of both soluble and insoluble Fe(III). Instead, genetic data suggests the triheme cytochromes PpcA-E, PgcA, and PpcA and PpcE homologs from *G. metallireducens* are promiscuous enough to support rapid and complete reduction of both soluble and insoluble Fe(III). As none of these cytochromes were required for electron transfer to electrodes, another as-yet unidentified periplasmic electron carrier is utilized during conductive biofilm growth.

The growth phenotypes of some deletion mutants differed from earlier insertional mutant data. For example, in prior washed cell U(VI) and Fe(III) reduction assays, Δ*ppcE*, Δ*ppcBC*, and Δ*ppcD* were reported to show slightly faster Fe(III) reduction (32). In addition, a comprehensive deletion of all five *ppc* paralogs eliminated *G. sulfurreducens*’ ability to reduce Fe(III) citrate (27). Along with variation expected from growth- vs. cell suspension assays, genetic factors could explain these differences. Tn-Seq data recently revealed many essential genes immediately up- and downstream of *ppc* paralogs that could have been affected by antibiotic cassette insertions, such as the cytochrome biosynthesis genes GSU0613-0614 adjacent to *ppcA*/GSU0612, DNA helicase GSU0363 downstream of *ppcCD*/GSU0364-365, and the purine metabolism cluster GSU1758-1759 adjacent to *ppcE*/GSU1760 (47). Improvements in genetic tools and genomic resequencing allowed use of verified markerless deletions to better avoid polar effects. Also, variations in expression between laboratory strains is common, especially for *pgcA* (47). A higher background level of PgcA likely aided the finding that this cytochrome can contribute to periplasmic electron transfer.

The discovery of a Δ*ppcABD* suppressor mutation in *pgcA* (*pgcA*^Δ37-39^) that rescued Fe(III) citrate growth (Fig 3) revealed an Ala^-2^-Gly^-1^-Cys^+1^ lipobox motif likely recognized by the *Geobacter* prolipoprotein diacylglyceryl transferase (Lgt) prior to cleavage by Lsp/SPII peptidase. This motif could be useful for targeting secretion of future heterologous proteins to the cell surface. It is interesting to note that PgcA was assigned a periplasmic localization in earlier proteomic studies (<u>p</u>eriplasmic <u>g</u>eobacter <u>c</u>ytochrome A), raising the possibility that a significant amount of this cytochrome is always present to the periplasm (48).

The ability of PgcA to aid Fe(III) reduction further underscored the promiscuity of periplasmic electron transfer (Fig 3 and Fig 4). While both are triheme *c*-type cytochromes, there is no significant amino acid sequence similarity between the 50 kDa PgcA and 10 kDa Ppc proteins, and PgcA contains long repetitive proline-threonine sequences between each heme. This raises the possibility that other outer membrane and extracellular multiheme cytochromes could participate in periplasmic electron transfer, explaining either the residual metal reduction activity in the Δ*ppc*5 Δ*pgcA* background or the growth phenotype of Δ*ppc*5 Δ*pgcA* mutants on poised electrodes.

With these new data, the question remains– why does *Geobacter* express multiple *ppc* paralogs at such high levels when such a metabolic burden appears unnecessary? A similar strategy, where different abundant periplasmic cytochromes appear to have overlapping functions, is also observed in the versatile metal-reducing bacterium *Shewanella oneidensis* (25). One hypothesis involves the ability of periplasmic cytochromes to act as ‘capacitors’, accepting electrons to enable constant proton motive force generation until extracellular oxidants can be found (25). Iron-starved *G. sulfurreducens* cells with fewer cytochromes have much slower rates of Fe(III) reduction, and cells subjected to on/off cycles of electrode polarization produce more net current while increasing cytochrome expression and electron storage capacity (49–51). Having multiple promiscuous carriers in the periplasm also increases the likelihood that new respiratory pathways can be acquired, as they could easily ‘plug in’ to the *Geobacter* network (22). The fact that *ppc* paralogs from other species fully complemented growth of mutants (Fig 7) indicates such horizontal exchange is feasible.

At every step of the *Geobacter* electron transfer chain, proteins that initially appeared redundant were later found to have non-overlapping roles. The inner membrane cytochromes ImcH and CbcL are both expressed constitutively, but only ImcH operates above ~0 V vs. SHE (10–13). The porin-cytochrome complexes OmcB and ExtABCD are both produced by cells in conductive biofilms, but only ExtABCD appears able to direct electrons to the electrode (14). Nanowire cytochromes OmcS and OmcE are linked to metal reduction, while only OmcZ is used for electrode reduction (16). The promiscuous Ppc family cytochromes show the opposite behavior, collecting electrons for distribution to any available acceptor, more similar to CctA and FccA in the *Shewanella* electron transfer network. Such versatility could greatly simplify full reconstruction of extracellular electron transfer in a heterologous host, and allow synthetic combinations of proteins from multiple species. In addition, the evidence that an undiscovered periplasmic carrier exists, which is only functional during electrode growth, provides a new target for engineering a separate communication network specifically for interaction with electrical surfaces.

## Materials and Methods

### Medium conditions and Inoculation

Strains and plasmids used in this study are listed in Table 1 and Table S1. Cloning information can be found in Table S2. *G. sulfurreducens* was grown in defined anaerobic salt medium with 20 mM acetate as the electron donor, and 40 mM fumarate, 55 mM Fe(III) citrate, or 30 mM amorphous Fe(III)-(oxyhydr)oxide as the acceptor as described (11, 38–40). The medium was prepared with 0.38 g/L KCl, 0.2 g/L NH4Cl, 0.069 g/L NaH_2_PO_4_·H_2_O, 0.04 g/L CaCl_2_, 0.2 g/L MgSO_4_·7H_2_O, 10 mL/L of a trace mineral mix, adjusted to pH to 6.8 and buffered with 2 g/L NaHCO_3_. The trace mineral mix contained 1.5 g/L nitrilotriacetic acid (NTA), 0.1 g/L MnCl_2_·_4_H_2_O, 0.5 g/L Fe_2_SO_4_·7H_2_O, 0.17 g/L CoCl_2_·6H_2_O, 0.10 g/L ZnCl_2_, 0.03 g/L CuSO_4_·5H_2_O, 0.005 g/L AlK(SO_4_)_2_·12H_2_O, 0.005 g/L H_3_BO_3_, 0.09 g/L Na_2_MoO_4_, 0.05 g/L NiCl_2_, 0.02 g/L NaWO_4_·2H_2_O, 0.10 g/L Na_2_SeO_4_. For media with Fe(III) citrate or Fe(III) oxide as the electron acceptor, the chelated trace mineral mix was replaced with non-chelated trace mineral mix which omitted NTA and instead dissolved minerals in 0.1 M HCl.

**TABLE 1.** Strains used in this study.

| Strains | Description or relative genotype | Source |
| --- | --- | --- |
| ***G. sulfurreducens*** |  |  |
| Wild type (WT) |  | Lab collection |
| DB1864 | $\Delta ppcA$ | This study |
| DB867 | $\Delta ppcB$ | This study |
| DB1483 | $\Delta ppcC$ | This study |
| DB1040 | $\Delta ppcD$ | This study |
| DB1854 | $\Delta ppcE$ | This study |
| DB1853 | $\Delta ppcBCDE$ ($ppcA^+$) | This study |
| DB1960 | $\Delta ppcACDE$ ($ppcB^+$) | This study |
| DB1917 | $\Delta ppcABDE$ ($ppcC^+$) | This study |
| DB1887 | $\Delta ppcABCE$ ($ppcD^+$) | This study |
| DB1850 | $\Delta ppcABCD$ ($ppcE^+$) | This study |
| DB1862 | $\Delta ppcABCDE$ ($\Delta ppc5$) | This study |
| DB1244 | $\Delta ppcABD$ | This study |
| DB1977 | $\Delta ppcABD$ $pgcA^{\Delta 37-39}$ | This study |
| DB1989 | $\Delta ppcABD$ $\Delta pgcA$ | This study |
| DB1546 | $\Delta pgcA$ | (37) |
| DB1976 | $\Delta ppc5$ $\Delta pgcA$ | This study |
| DB1900 | $\Delta ppc5$ Tn7::$ppcA$ | This study |
| DB2161 | $\Delta ppc5$ Tn7::$ppcC$ | This study |
| DB2173 | $\Delta ppc5$ Tn7::$ppcA_{SP}$-$ppcE$ | This study |
| DB2005 | $\Delta ppc5$ $\Delta pgcA$ Tn7::$ppcA$ | This study |
| DB2064 | $\Delta ppc5$ $\Delta pgcA$ Tn7::$ppcC$ | This study |
| DB2007 | $\Delta ppc5$ $\Delta pgcA$ Tn7::$ppcE$ | This study |
| DB2062 | $\Delta ppc5$ $\Delta pgcA$ Tn7::$ppcA_{SP}$-$ppcE$ | This study |
| DB2148 | $\Delta ppc5$ $\Delta pgcA$ Tn7::$ppcE_{SP}$-$ppcA$ | This study |
| DB2149 | $\Delta ppc5$ $\Delta pgcA$ Tn7::$ppcA_{SP}$-$Gmet$ $ppcA$ | This study |
| DB2210 | $\Delta ppc5$ $\Delta pgcA$ Tn7::$ppcA_{SP}$-$Gmet$ $ppcE$ | This study |
| DB2150 | $\Delta ppc5$ $\Delta pgcA$ Tn7::$ppcA_{SP}$-DVU3171 | This study |
| DB2063 | $\Delta ppc5$ $\Delta pgcA$ Tn7::$ppcA_{SP}$-$cctA$ | This study |
| ***E. coli*** |  |  |
| UQ950 | Cloning strain |  |
| S17-1 | Conjugation donor strain |  |
| MFDpir | Conjugation donor strain |  |
| DB1325 | MFDpir conjugation donor strain containing helper plasmid pmobile-CRISPRi | (44) |
|  | UQ950 containing pRK2-Geo2-lacZa | (38) |
| DB1777 | UQ950 containing pTn7m-kan-lacZa | This study |
| DB1880 | UQ950 containing pTn7-Geo7 | (17) |
| DB1284 | S17-1 containing pDppcA | This study |
| DB1283 | S17-1 containing pDppcB | This study |
| DB968 | S17-1 containing pDppcC | This study |
| DB1003 | S17-1 containing pD*ppcBC* | This study |
| DB943 | S17-1 containing pD*ppcD* | This study |
| DB1851 | S17-1 containing pD*ppcE* | This study |
| DB1782 | S17-1 containing pD*pgcA* | This study |
| DB1927 | S17-1 containing pD*pgcA* wo1760 | This study |
| DB1914 | S17-1 containing pGeo7::*ppcA* | This study |
| DB2056 | S17-1 containing pGeo7::*ppcC* | This study |
| DB1945 | S17-1 containing pGeo7::*ppcE* | This study |
| DB2058 | S17-1 containing pGeo7::*ppcA*<sub>SP</sub>-*ppcE* | This study |
| DB2133 | S17-1 containing pGeo7::*ppcE*<sub>SP</sub>-*ppcA* | This study |
| DB2134 | S17-1 containing pGeo7::*ppcA*<sub>SP</sub>-*Gmet ppcA* | This study |
| DB2144 | S17-1 containing pGeo7::*ppcA*<sub>SP</sub>-*Gmet ppcE* | This study |
| DB2135 | S17-1 containing pGeo7::*ppcA*<sub>SP</sub>-DVU3171 | This study |
| DB2192 | MFDpir containing pGeo7::*ppcA*<sub>SP</sub>-*cctA* | This study |

To make Fe(III) oxide, 10 g of FeSO_4_·7H_2_O was added to 1 L of water, and 5.32 mL of 30 % H_2_O_2_ added with stirring overnight. This produced schwertmannite (Fe_8_O_8_(OH)_6_(SO_4_)·nH_2_O), which was washed in distilled water and stored until needed. After addition of schwertmannite to medium and autoclaving, the Fe(III) ages to an amorphous Fe(III)-(oxyhydr)oxide, allowing generation of a highly repeatable iron oxide form between experimental replicates.

For electrode bioreactor media, 40 mM acetate was added as the electron donor and a poised graphite electrode (+0.24 V vs. SHE) used as the acceptor. 50 mM NaCl was added for osmotic balance to replace salts present in typical fumarate or Fe(III)-citrate growth medium. All media were flushed with N_2_:CO_2_ (80:20) gas mix passed through a heated copper column to remove oxygen.

All experiments were initiated by streaking out frozen stocks onto anaerobic 1.7 % agar medium containing 20 mM acetate as the electron donor and 40 mM fumarate as the electron acceptor in MACS MG-500 gloveless anaerobic chamber, (Don Whitley Scientific) with N_2_:CO_2_:H_2_ (75:20:5). Trypticase peptone (0.1%) and cysteine (1 mM) were added to promote recovery on the solid medium. Single colonies were picked and propagated in triplicate 1 mL liquid medium with 20 mM acetate and 40 mM fumarate, inoculated 1:10 into 10 mL medium for use in experiments. For metal reduction experiments, cultures at 0.6 OD_600_ were inoculated 1:100 into medium containing 20 mM acetate as the electron donor and 55 mM Fe(III) citrate or 30 mM Fe(III) oxide as the electron acceptor. All cultures were incubated at 30 °C.

### Fe(III) reduction assay

Fe(III) citrate and Fe(III) oxide medium samples were diluted 1:10 into 0.5 N HCl for each timepoint. The solution was diluted further with 0.5 N HCl when needed. Samples were analyzed for Fe(II) using a modified FerroZine assay (41). The FerroZine solution contained 2 g/L FerroZine and 23.8 g/L HEPES with pH adjusted to 7.0. 300 μl of the FerroZine solution was added to 50 μl of the diluted samples in 96 well plates. The plates were read at 625 nm by BioTek Synergy multi-mode reader.

### Growth with poised electrode as electron acceptor

Three-electrode bioreactors consist of a 3 cm^2^ graphite working electrode set at +0.24 V vs. SHE, a platinum counter electrode, and a calomel reference electrode. The graphite working electrode was polished with P1500 sandpaper and sonicated before each use. Each bioreactor was inoculated with 12 mL of medium with 40 mM acetate and 50 mM NaCl, and flushed with humidified N_2_:CO_2_ (80:20) gas overnight, before inoculation of 4 mL OD_600_ 0.5 cells. Cells were grown in 30°C under constant stirring. Biomass attached to anodes was determined by removing electrodes during exponential growth (as they reached 100 μA/cm^2^), boiling in 0.2 N NaOH, and determining the total protein concentration using the Bicinchoninic acid (BCA) assay.

### Gene deletion and complementation

A sucrose-SacB counter-selection method was employed for construction of scarless deletion strains (38). Up- and downstream fragments (~ 750 bp each) of the target gene were joined by overlapping PCR and ligated into pK18*mobsacB*. The plasmid was purified and Sanger-sequenced to verify the target region after transformation into *E. coli* UQ950. Once confirmed, the plasmid was transformed into *E. coli* S17-1 or MFDpir to be conjugated with *G. sulfurreducens.* 1 mL of each *G. sulfurreducens* recipient strain and S17-1 or MFDpir donor strain culture were centrifuged together, decanted, and resuspended in the residual supernatant. This cell suspension was placed on sterilized 0.22 um pore size filter paper on agar medium with 20 mM acetate and 40 mM fumarate overnight. Merodiploids were selected on agar medium containing 20 mM acetate and 40 mM fumarate with 200 μg/mL kanamycin, and integration of the plasmid at the target site verified by PCR. Colonies with the integrated *sacB-*containing plasmid were subjected to sucrose-counter selection on solid agar medium containing 20 mM and 40 mM fumarate with 10% sucrose, to screen for recombination of homologous regions which should delete the target gene in 50% of events. PCR of re-isolated antibiotic-sensitive colonies using flanking primers was performed to identify deletion strains.

### Suppressor analysis

Replicate cultures of mutants with significant Fe(III) citrate reduction defects, (Δ*ppcABD*, Δ*ppcABCDE*, Δ*ppcABCDE*Δ*pgcA*) were grown with 20 mM acetate and 40 mM fumarate, then inoculated 1:100 into medium with 20 mM acetate and 55 mM Fe(III) citrate. If growth was detected, cultures were subcultured with Fe(III) citrate, and tubes demonstrating growth faster than parent cultures then streaked onto plates of agar medium containing 20 mM acetate and 40 mM fumarate to isolate colonies. Individual colonies were rescreened in medium with 20 mM acetate and 55 mM Fe(III) citrate to identify clonal suppressor strains. Genomic DNA of these strains was resequenced along with the parent, and breseq version 0.28.0 used to identify mutations compared to the parent.

### Fractionation of periplasmic proteins

A protocol for releasing periplasmic proteins via osmotic shock was adapted from (42). Periplasmic fractions for Fig 1C, 2C, 3D, 4C, 5C, 5E, 5G, 6B, 7B, S1, S4, S5B, and S6B are from stationary phase fumarate-grown cultures. Cultures were harvested and adjusted by OD_600_ so all extractions began with the same amount of cells. Fig S2 shows periplasmic fractions from Fe(III) citrate-grown cultures that reduced Fe(III) to ~ 50 mM.

Cultures were centrifuged at 5,200 x g for 10 minutes, pellets were resuspended in 1 mL 50 mM Tris, 250 mM sucrose, 2.5 mM EDTA, pH 8.0, and equilibrated at room temperature for 5 minutes. This suspension was centrifuged at 16,000 x g at 4 °C for 10 minutes and the supernatant carefully removed. The pellet was rapidly resuspended in with 200 μl of ice-cold 5 mM MgSO_4_ with gentle mixing on ice, causing rapid influx of water to the periplasm, osmotically disrupting the EDTA-destabilized outer membrane, and releasing periplasmic proteins. After 30 minutes, cells and debris were removed by centrifugation at 16,000 x g for 10 minutes at 4 °C. For electrode-grown cells, biofilms were collected by rinsing off four electrodes with a pipette tip in a tube containing 500 μl of 50 mM Tris, 250 mM sucrose, 2.5 mM EDTA, pH 8.0 and resuspended in 5 mM 300 μl of MgSO_4_ solution. 15 mL of planktonic cultures were collected in 1 mL of 50 mM Tris, 250 mM sucrose, 2.5 mM EDTA, pH 8.0 and resuspended in 280 μl of MgSO_4_ solution. Electrode-grown cells were collected at a stationary phase.

The supernatant containing the periplasmic fraction was boiled at 95 °C for 5 minutes with SDS loading buffer (omitting β-Mercaptoethanol) and separated on a tricine-SDS-PAGE gel (adapted from (43)). The 16% resolving gels were made from the solution containing 5.33 mL acrylamide/bis-acrylamide 19:1 30% (w/v), 4.3 mL of 2.5 M Tris buffer (pH 8.8), 0.22 mL of dH_2_O with 100 μl of 30 mg/mL APS and 6 μl of TEMED added to polymerize. The 7% resolving gels were made from 2.33 mL acrylamide/bis-acrylamide 19:1 30% (w/v), 5.6 mL of 2.5 M Tris buffer (pH 8.8), 1.91 mL of dH_2_O with 150 μl of 30 mg/mL APS and 7 μl of TEMED. The stacking gels were made from the solution containing 0.66 mL acrylamide/bis-acrylamide 19:1 30% (w/v) solution, 0.76 mL 2.5 M Tris buffer (pH 8.8), 3.42 mL of dH_2_O with 150 μl APS 30 mg/mL and 5 μl TEMED. All heme-staining periplasmic fraction gels are 16% tricine gels, except Fig 3D and S2A. For detection of *c*-type cytochromes, the gel was dark-incubated for 1 hour in a solution containing 0.0227 g of 3,3’,5,5’-Tetramethylbenzidine dissolved in 15 mL of methanol and mixed with 35 mL 0.25 M sodium acetate (pH 5.0) for a total 50 mL. The gel was visualized upon the addition of 1.5 mL of 3 % H_2_O_2_ for 10-15 minutes.

To test if periplasmic fractions contained contaminating membranes, we performed ultracentrifugation of periplasmic fractions at 177,000 g at 4 °C for 45 minutes (Fig S1). The samples of pre- and post-ultracentrifugation did not show any significant difference.

### Hydrogen Peroxide tolerance assay

An assay for hydrogen peroxide (H_2_O_2_) sensitivity was conducted using an H_2_O_2_ disc diffusion assay. 10 mL media containing melted 0.5% molten agar (‘0.5% top agar’) was mixed 1:100 from liquid culture of OD_600_ 0.6 was poured on top of the solid medium containing 1.7% agar and 0.1% Trypticase. After 3 hours, autoclaved BBL Blank Paper Discs (6 mm diameter) were placed on top agar and spotted with 10 μl of 400 mM H_2_O_2_. The diameter of the zone of inhibition was measured after 2.5 days of incubation at 30 °C in MACS MG-500 gloveless anaerobic chamber, Don Whitley Scientific. The zone of inhibition around each filter disc was measured by ImageJ.

## Acknowledgements

SC and DB were supported by the Office of Naval Research (Award Number N00014-18-1-2632). We thank Jeff Gralnick for helpful discussions.

## Supplementary Materials

**Fig S1.**
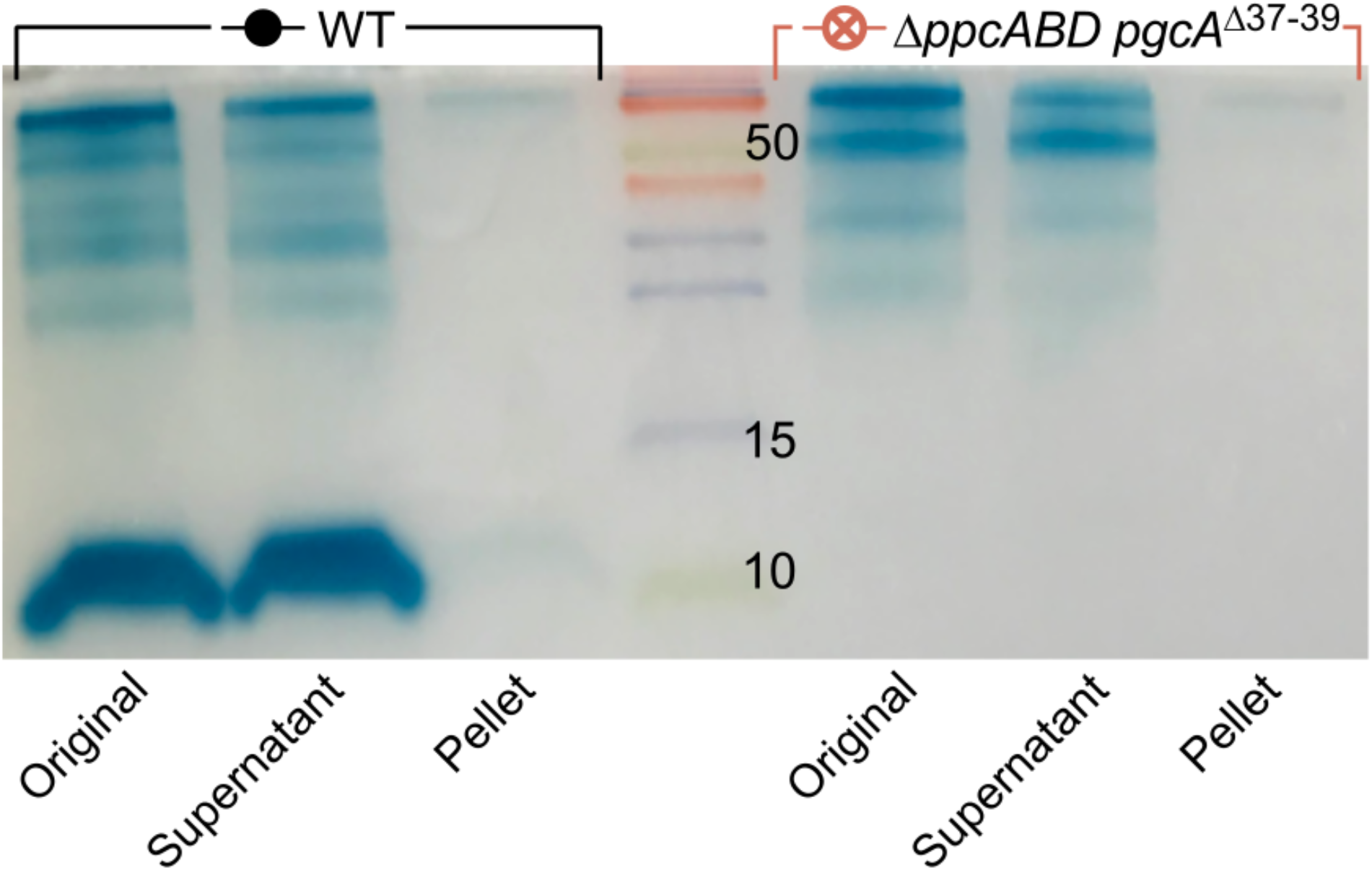
Periplasmic fractions pre- and post-ultracentrifugation. To test if periplasmic fractions contained significant membrane-bound cytochromes, samples were subjected to 177,000 g for 45 min. **Original**: total periplasmic fraction before ultracentrifugation. **Supernatant**: supernatant after ultracentrifugation. **Pellet**: resuspended fraction of the ultracentrifugation pellet.

**Fig S2.**
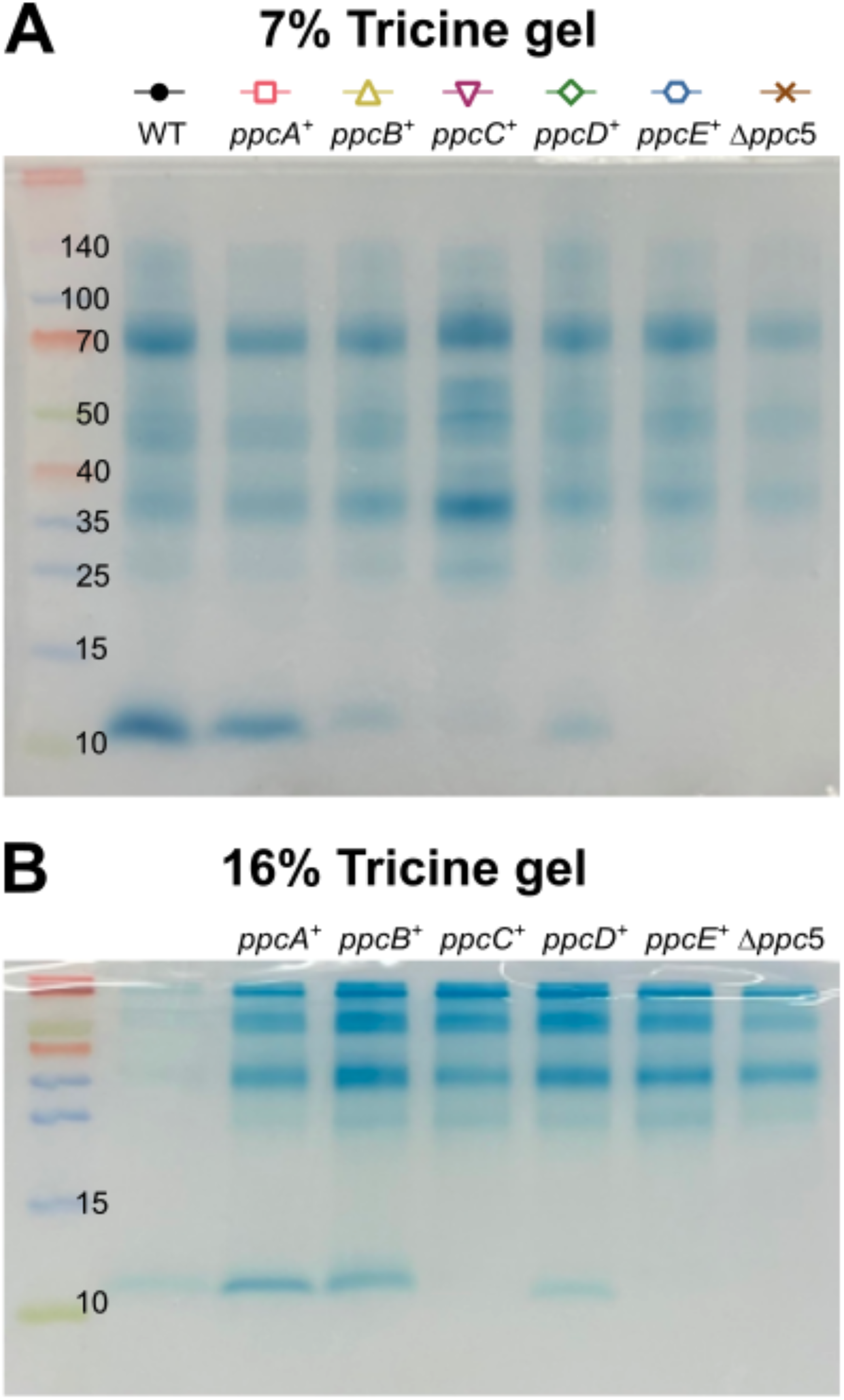
Periplasmic cytochrome fractions from Fe(III) citrate-grown cultures showing similar cytochrome abundance as with fumarate grown cells. (A) 7% tricine gel. (B) 16% tricine gel.

**Fig S3.**
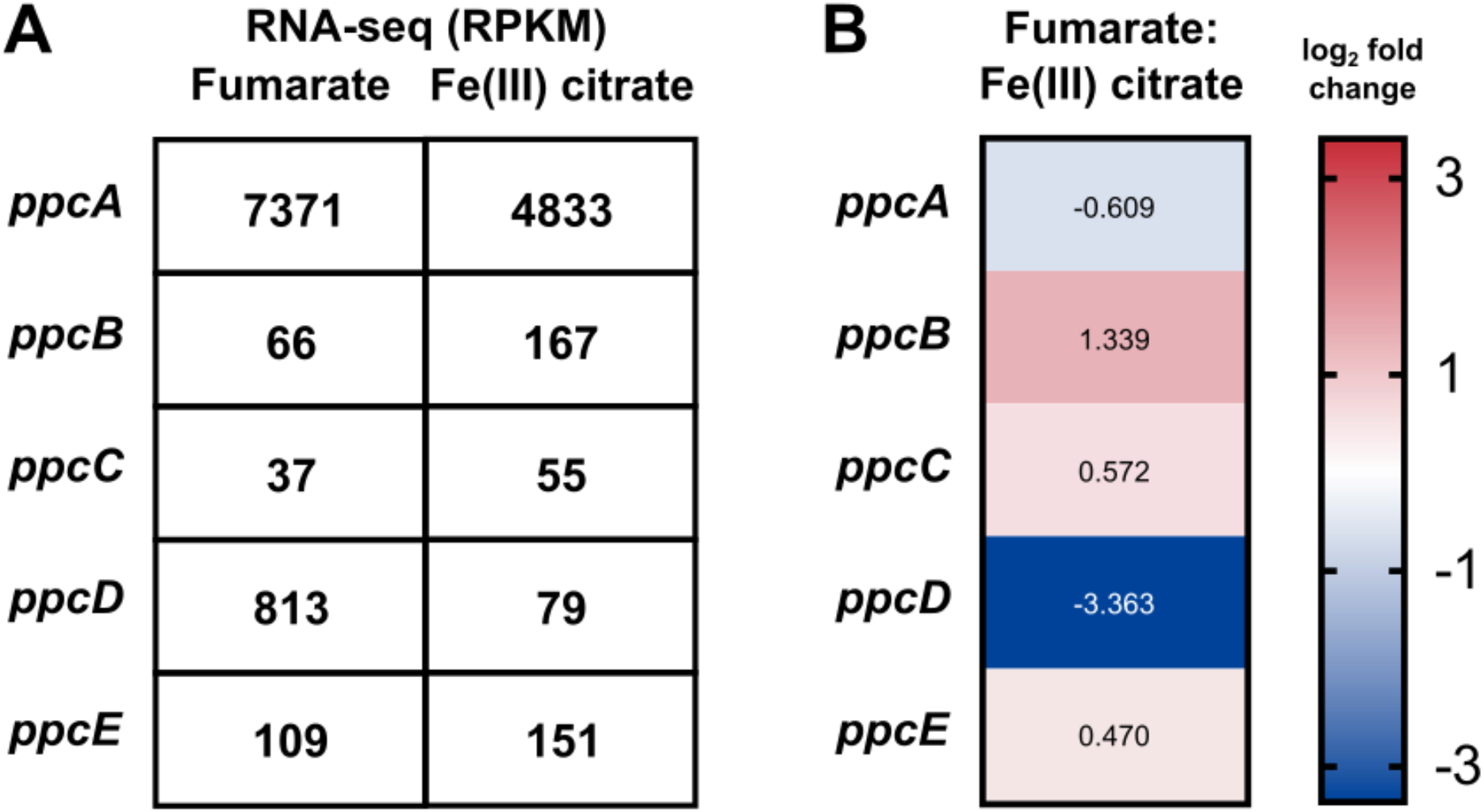
Expression level of Ppc paralog genes comparing fumarate to Fe(III) citrate cultures *G. sulfurreducens.* Large expression differences have been reported, but primarily from stationary phase cultures or other genetic backgrounds. Adapted from RNA-seq data from Joshi et al. 2020. (A) RNA-seq RPKM values of Ppc-family cytochrome transcripts in *G. sulfurreducens.* (B) Log2 fold expression change from fumarate to Fe(III) citrate culture.

**Fig S4.**
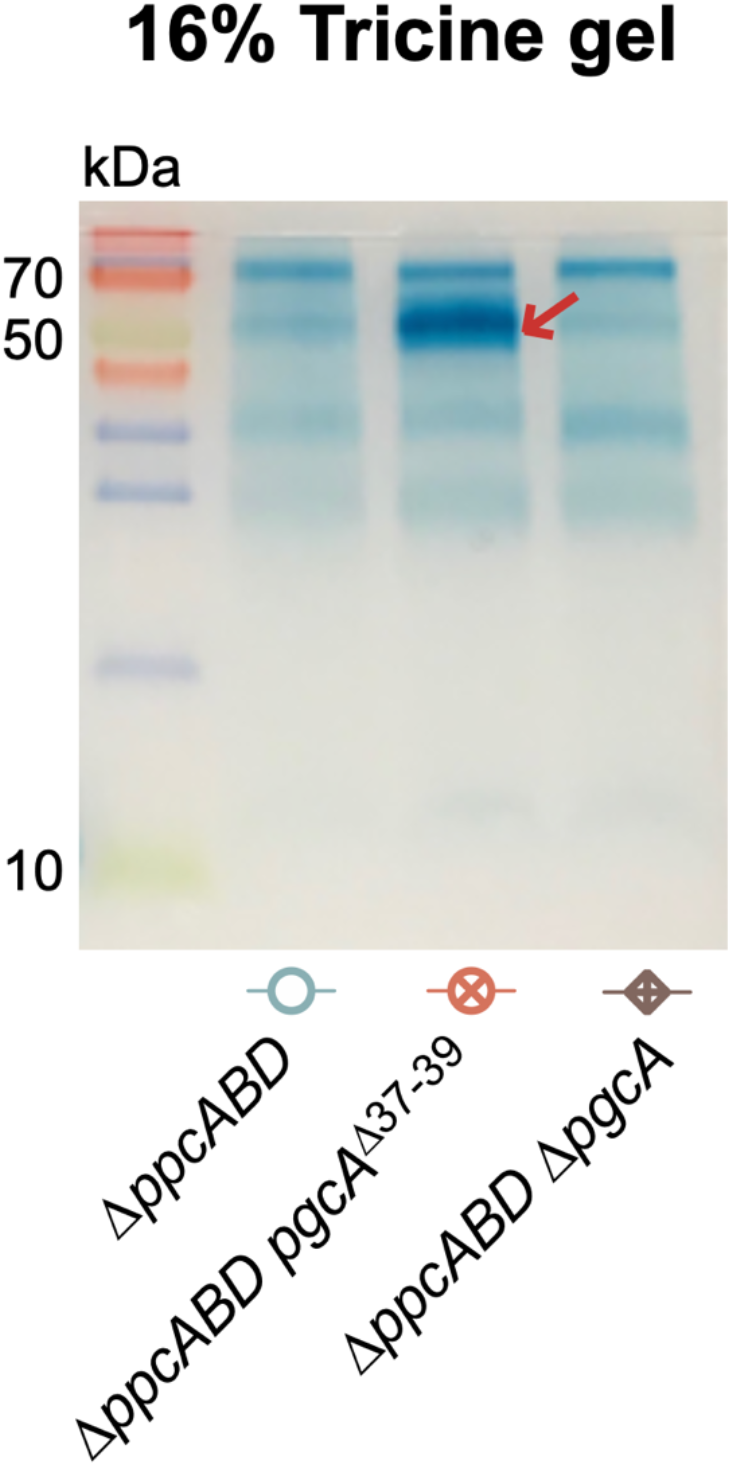
Heme-staining of the samples from Fig 3D on a 16% tricine gel. Proteins with molecular weight over 20 kDa are shown using 16% tricine gel as comparison.

**Fig S5.**
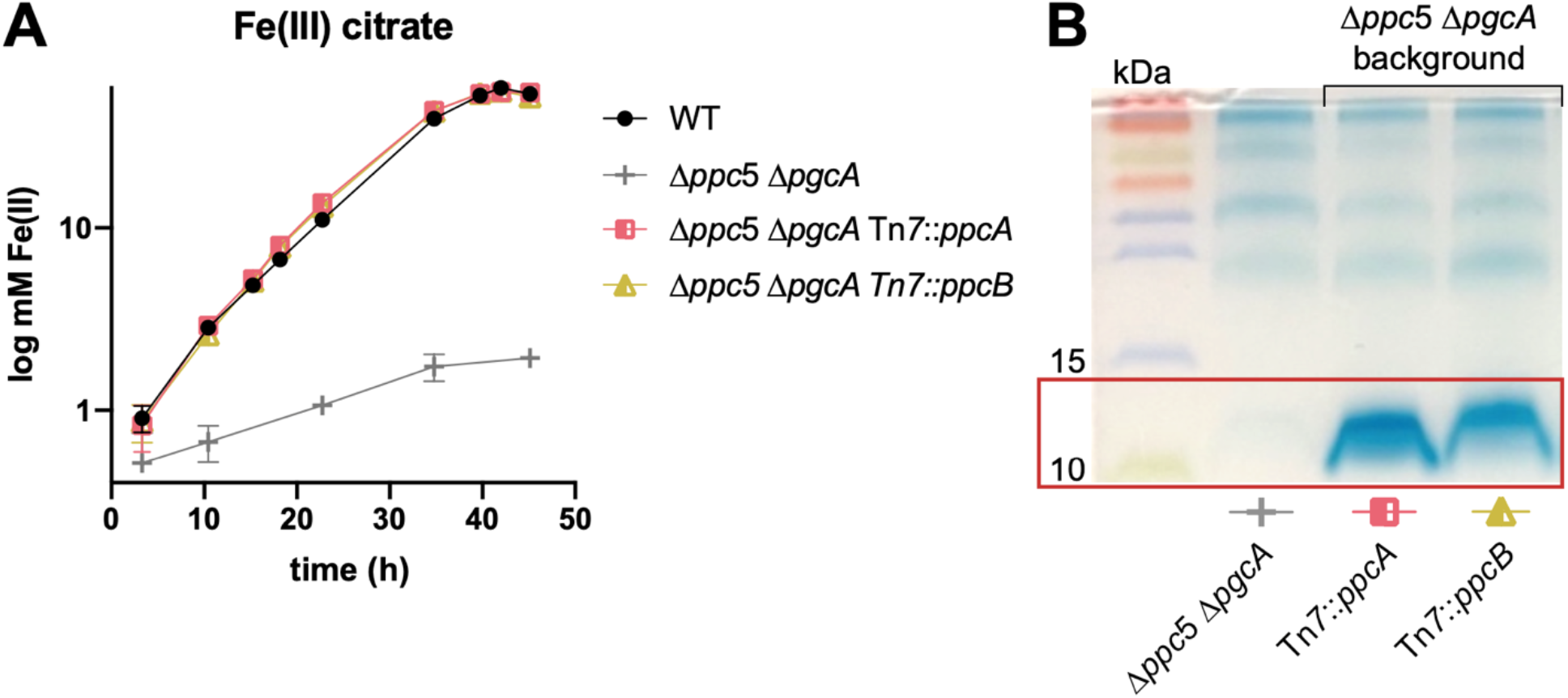
Re-introduction of *ppcB* rescues wild type level Fe(III) citrate reduction in Δ*ppcABCDE*Δ*pgcA*. (A) Fe(III) citrate reduction. (B) Heme-staining of periplasmic fraction. Results were similar to strain containing only PpcB in its native context (see Fig 2).

**Fig S6.**
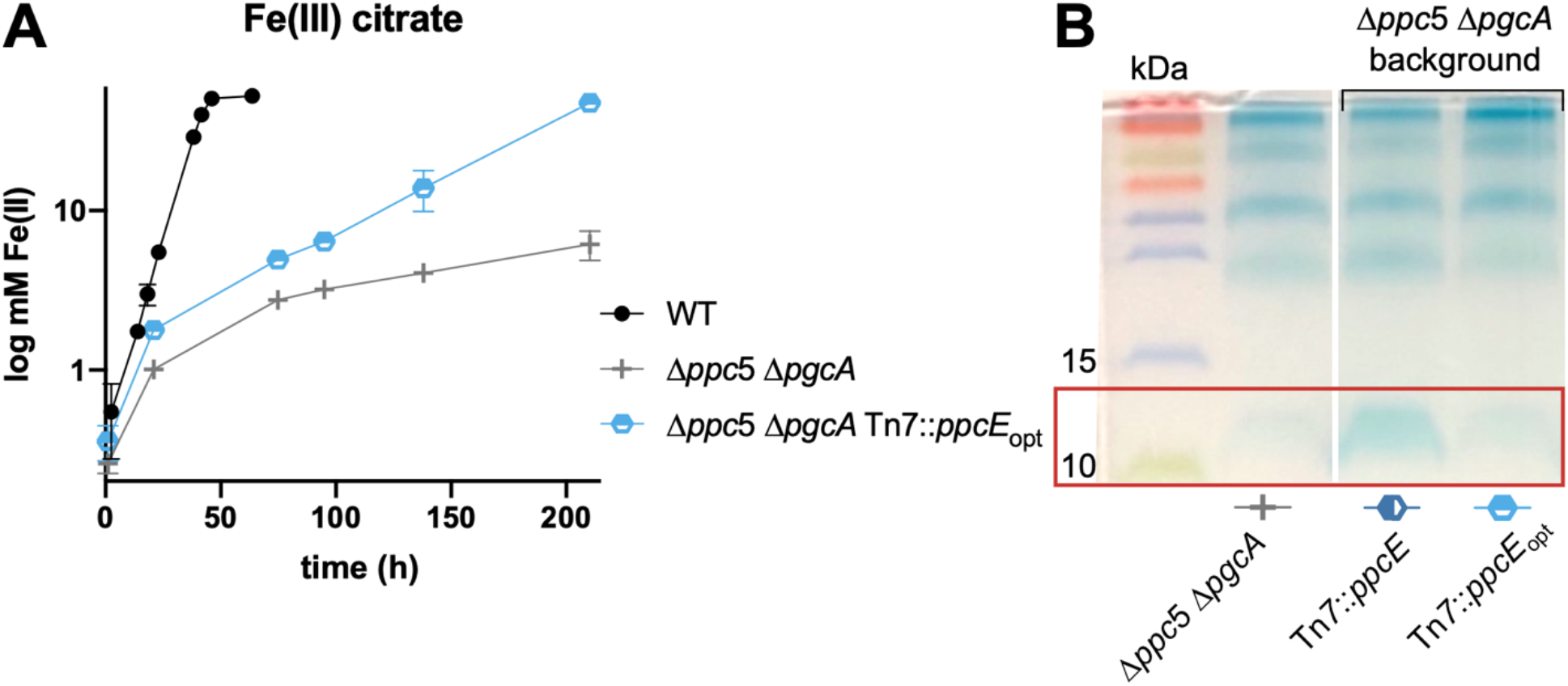
Codon-optimization does not increase the abundance of PpcE in periplasm. A separate experiment was conducted to determine if codon optimization could increase PpcE abundance (A) Fe(III) citrate reduction. (B) Heme-staining of periplasmic fraction. This is the part of the gel picture removed from Fig 5C, as this line of investigation was not pursued further.

**Fig S7.**
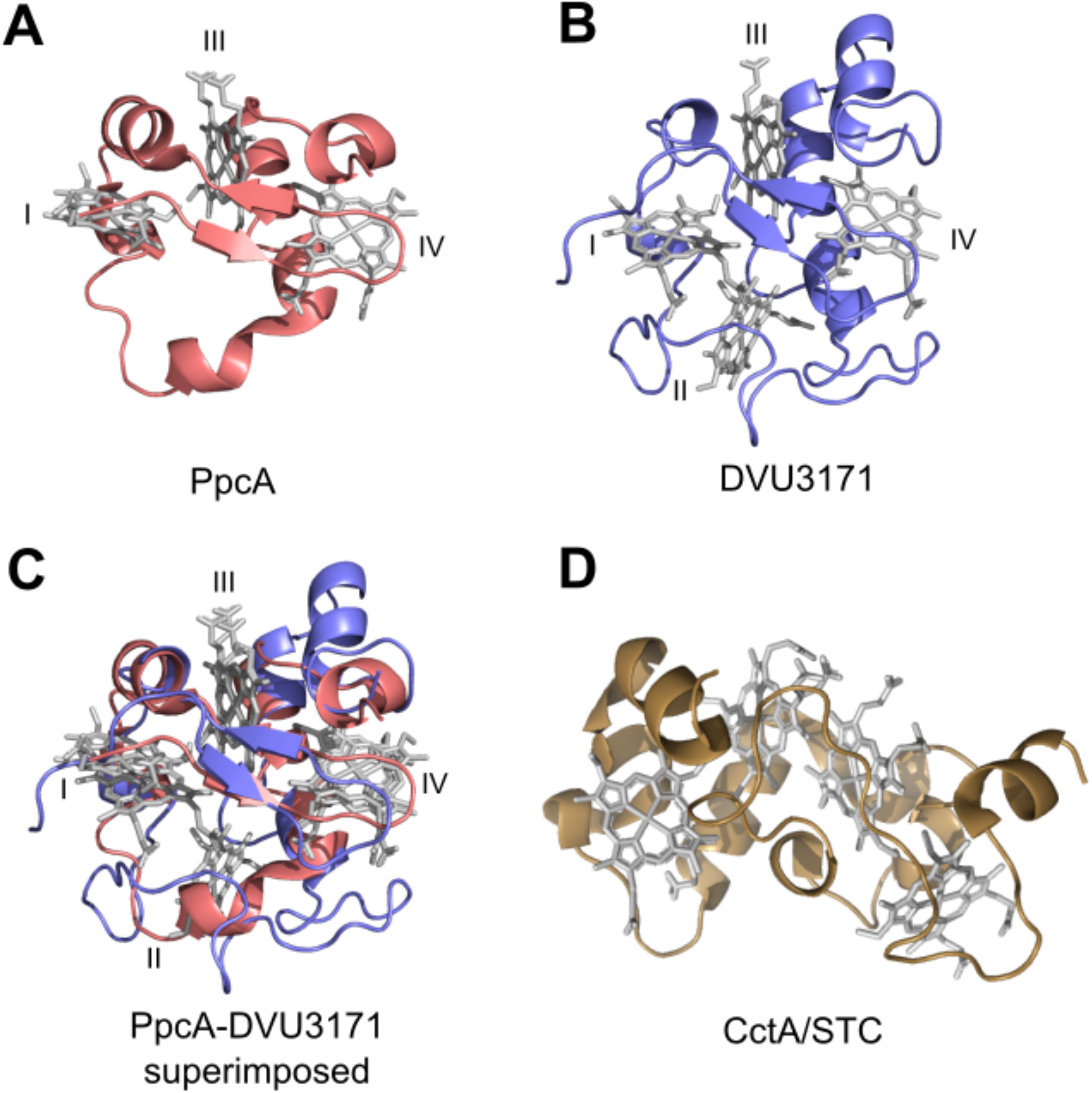
Structures of periplasmic cytochromes expressed in this work. Hemes are shown in gray, designated as heme I, II, III, or IV. (A) PpcA from *G. sulfurreducens* (PDB: 1OS6). (B) DVU3171 from *D. vulguris* (Hildenborough) (PDB: 2CTH) (Simoes et al., 1998) that has an additional heme group (heme II). (C) Structures of PpcA and DVU3171 superimposed on PyMOL. (D) CctA/STC from *S. oneidensis* (PDB: 1M1Q) (Leys et al., 2002).

**Fig S8.**
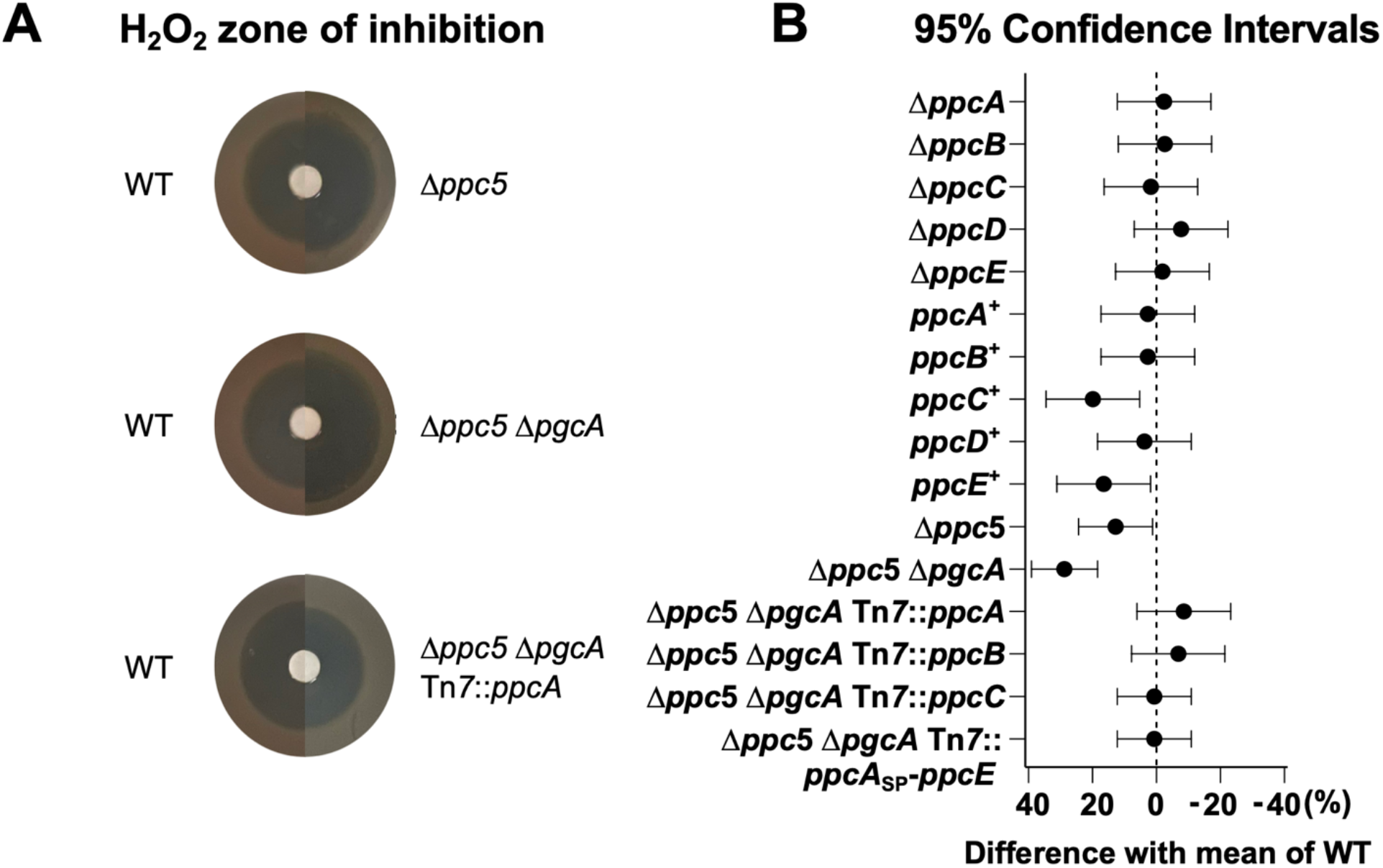
H_2_O_2_ sensitivity of periplasmic cytochrome mutants. (A) Pictures showing examples of inhibition zone. (B) Area of inhibition for each mutant compared with WT (Dunnett’s multiple comparison test). Strains where periplasmic fractions show little to no 10 kDa cytochrome (*ppcC*^+^, *ppcE^+^,* quintuple, and sextuple mutants) show slightly reduced H_2_O_2_ tolerance. Each value is calculated by subtracting the WT value from each mutant. A more positive difference compared to the WT mean (points shifted left) indicates increased sensitivity to oxidative stress, and a larger zone of inhibition by the mutant.

**Table S1.** Plasmids used in this study.

| Plasmid | Insert | Restriction Enzyme |
| --- | --- | --- |
| **pK18mobsacB** |  |  |
| pDppcA | iDppcA | BamHI-SacI |
| pDppcB | iDppcB | BamHI-SacI |
| pDppcC | iDppcC | HindIII-EcoRI |
| pDppcBC | iDppcBC | HindIII-EcoRI |
| pDppcD | iDppcD | HindIII-EcoRI |
| pDppcE | iDppcE | BamHI-EcoRI |
| pDpgcA | iDpgcA | BamHI-SacI |
| pDpgcA_wo1760 | iDpgcA_wo1760 | BamHI-EcoRI |
| **pRK2-Geo2-lacZa** |  |  |
| **pTn7m-kan-lacZa** | ikan-lacZa |  |
| **pTn7-Geo7** | iGeo7 | AscI-PmeI |
| pGeo7::ppcA | iGeo7-ppcA | NdeI-BamHI |
| pGeo7::ppcC | iGeo7-ppcC | NdeI-ApaLI |
| pGeo7::ppcE | iGeo7-ppcE | NdeI-BamHI |
| pGeo7::ppcA<sub>SP</sub>-ppcE | iGeo7-ppcA<sub>SP</sub>-ppcE | NdeI-ApaLI |
| pGeo7::ppcE<sub>SP</sub>-ppcA | iGeo7-ppcE<sub>SP</sub>-ppcA | NdeI-ApaLI |
| pGeo7::ppcA<sub>SP</sub>-Gmet ppcA | iGeo7-ppcA<sub>SP</sub>-Gmet ppcA | NdeI-ApaLI |
| pGeo7::ppcA<sub>SP</sub>-Gmet ppcE | iGeo7-ppcA<sub>SP</sub>-Gmet ppcE | NdeI-ApaLI |
| pGeo7::ppcA<sub>SP</sub>-DVU3171 | iGeo7-ppcA<sub>SP</sub>-DVU3171 | NdeI-ApaLI |
| pGeo7::ppcA<sub>SP</sub>-cctA | iGeo7-ppcA<sub>SP</sub>-cctA | NdeI-ApaLI |

**Table S2.**
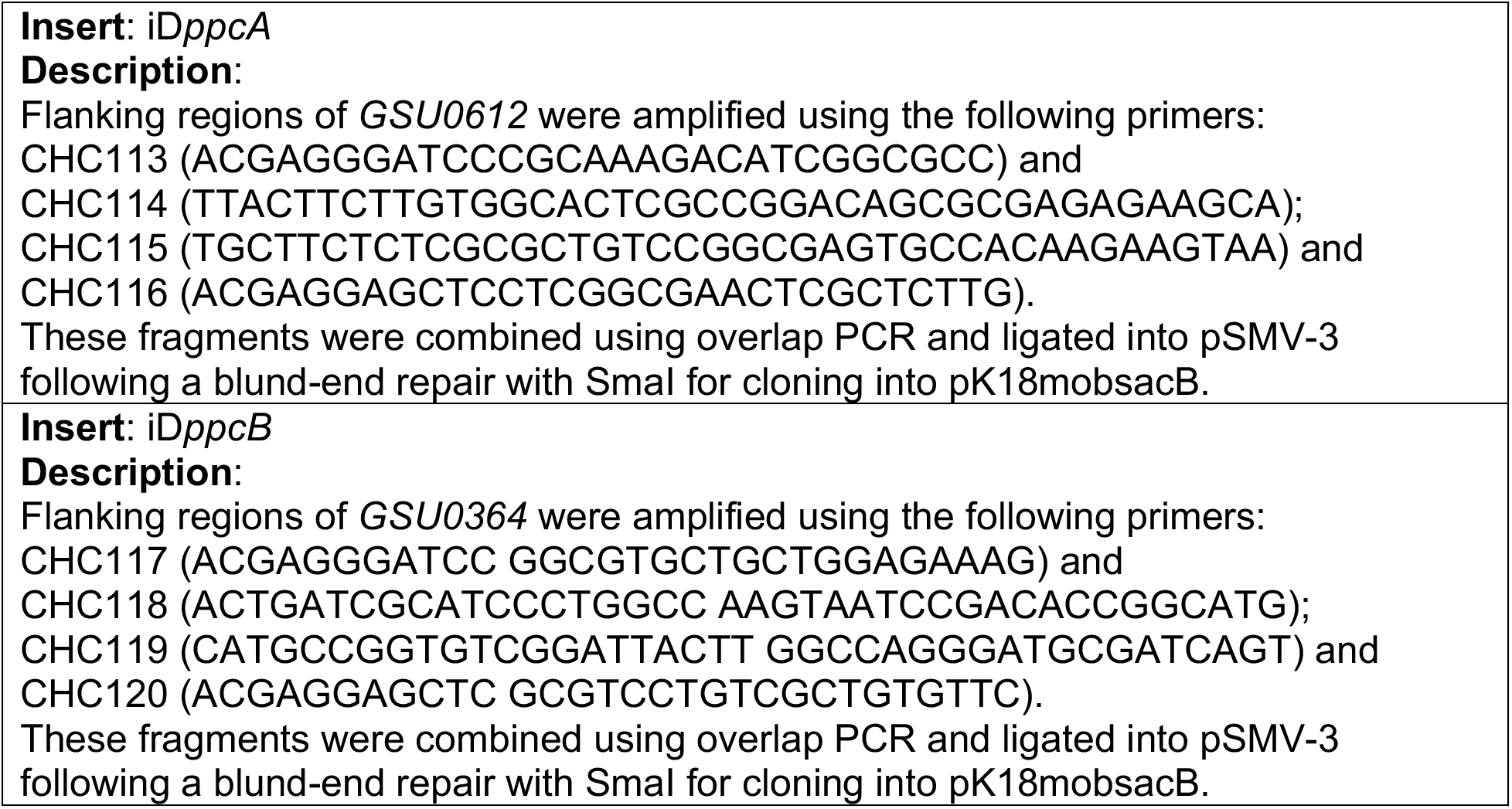
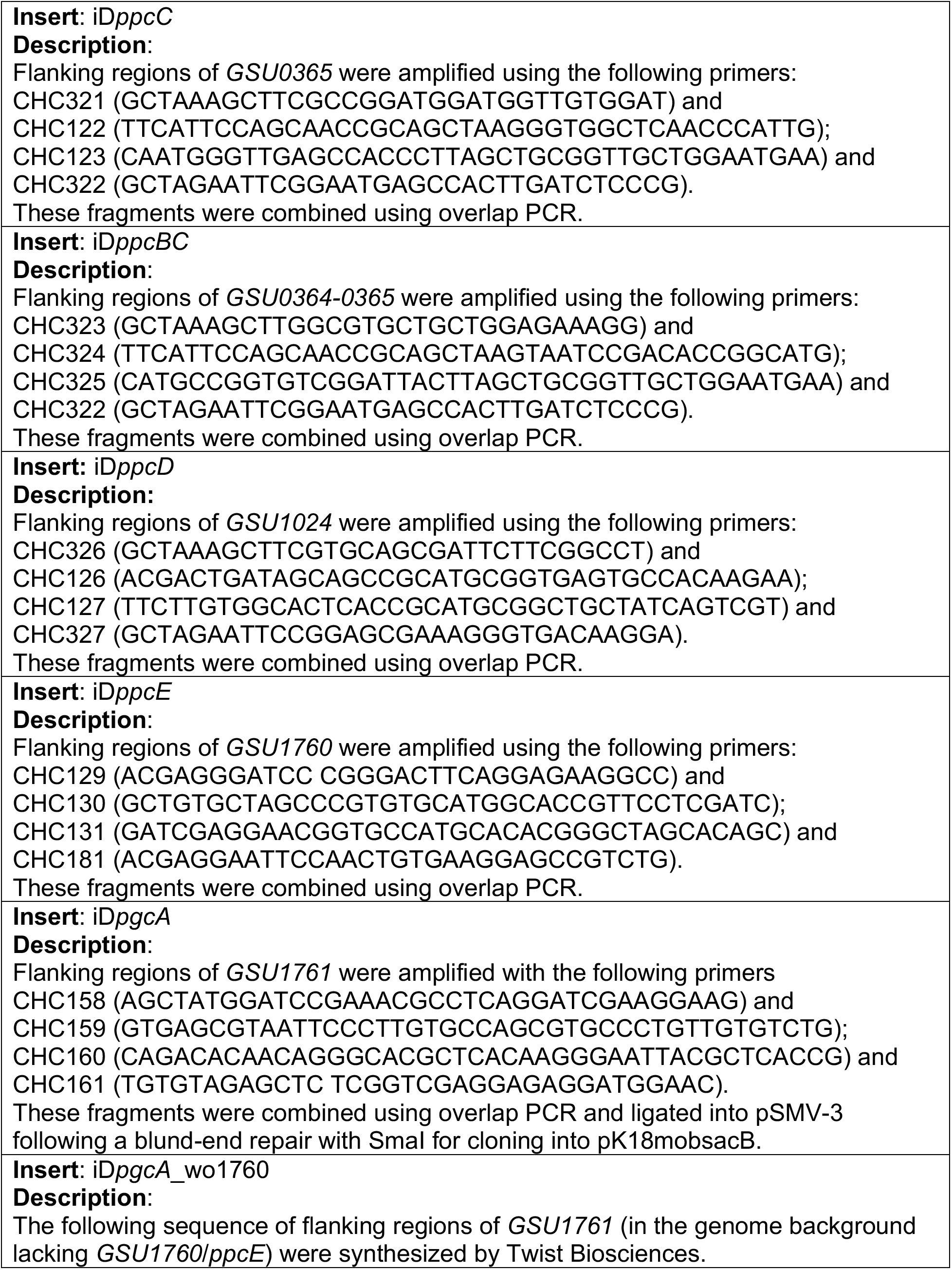
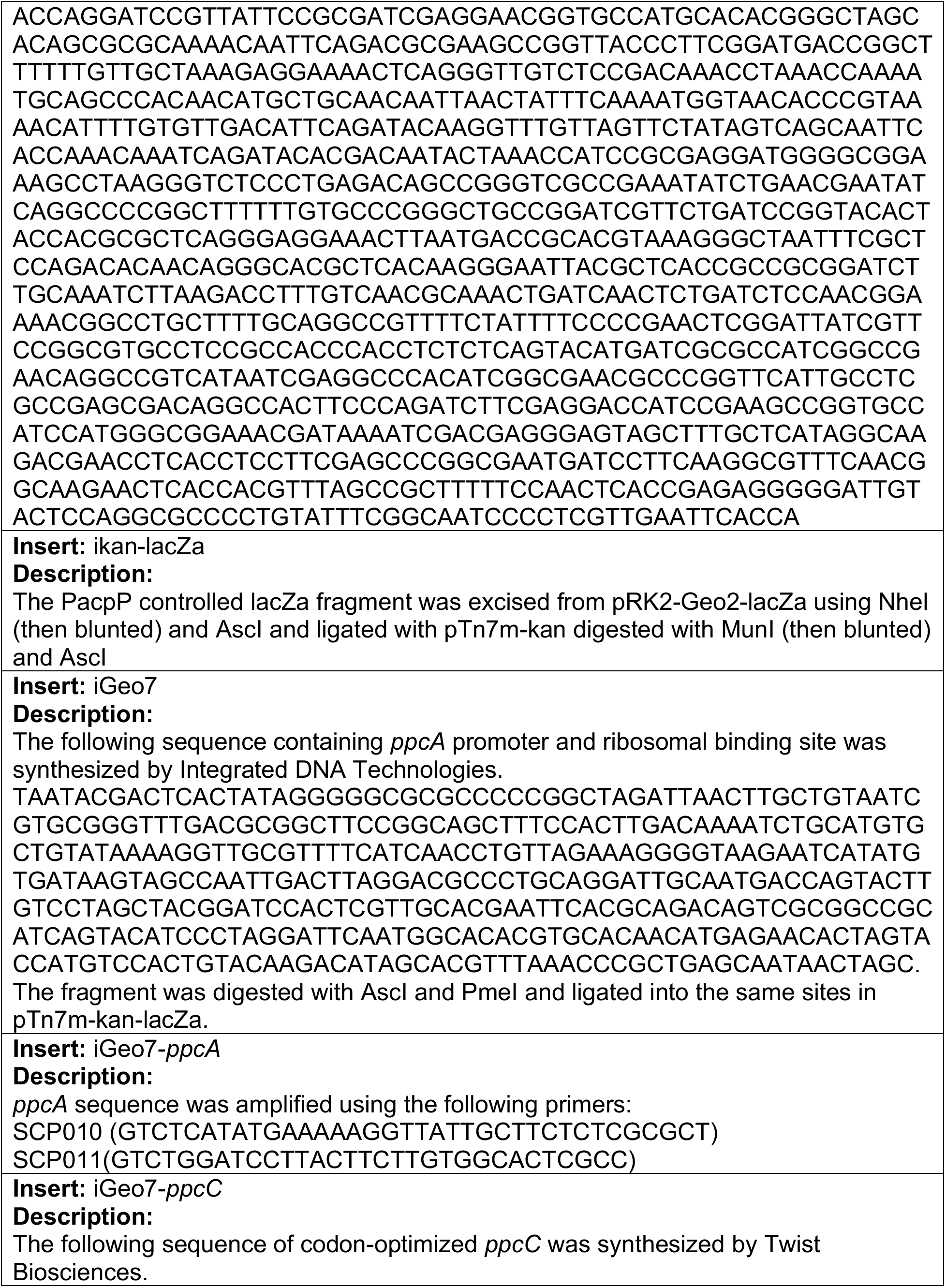
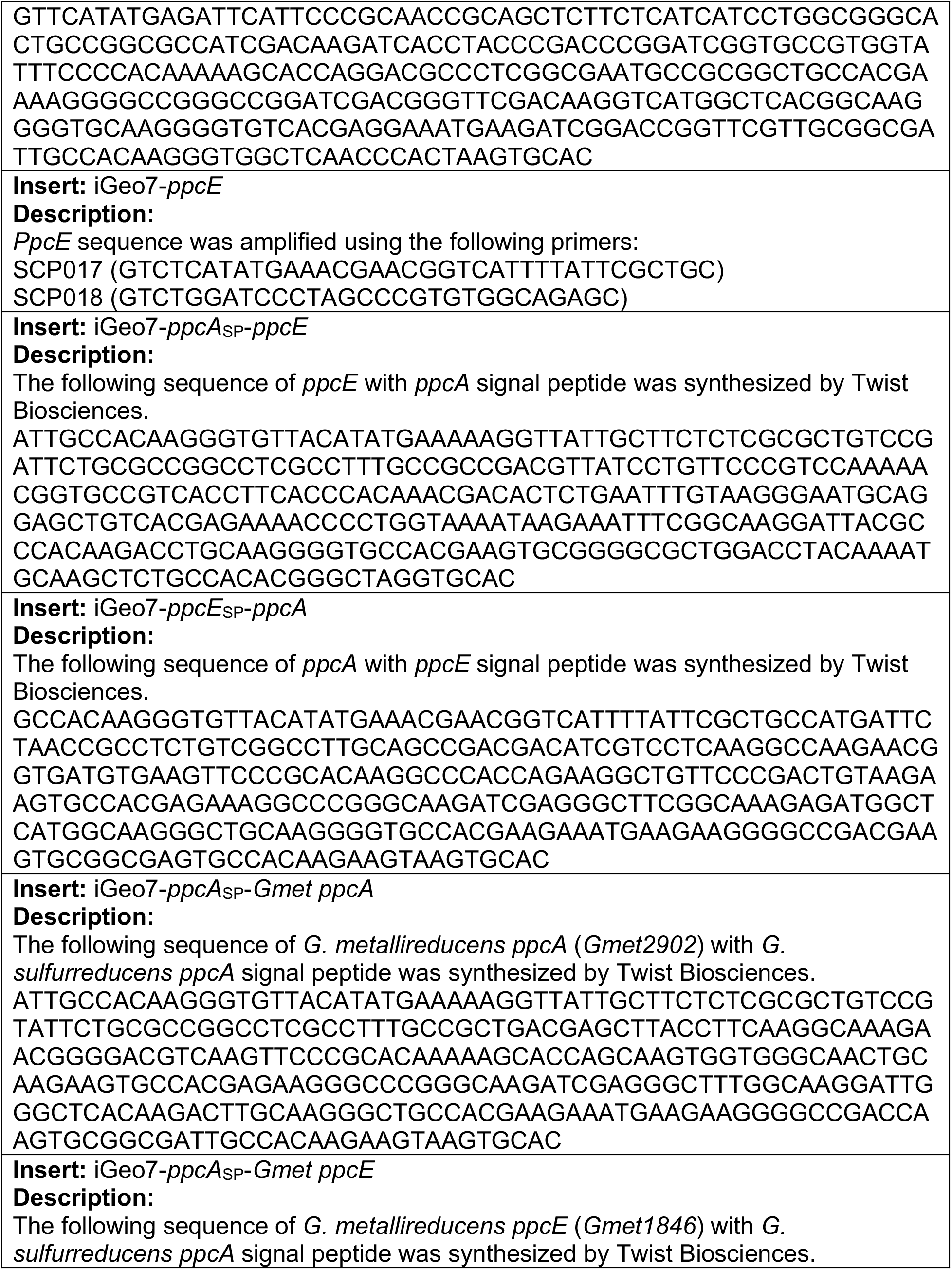
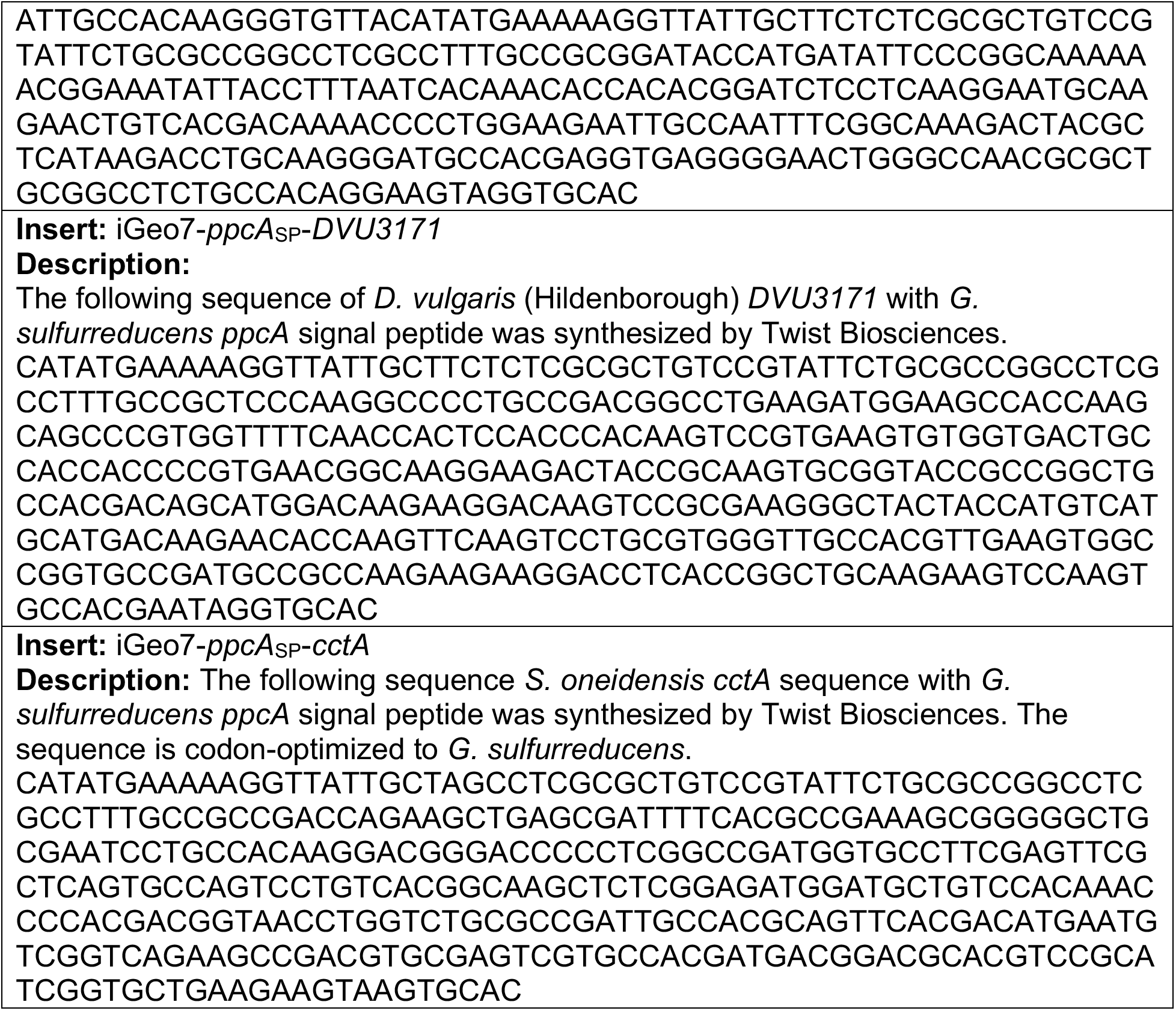
Insert information for cloning.

## References Cited

1. Gralnick JA, Newman DK. 2007. Extracellular respiration. Molecular Microbiology 65:1–11.

2. Edwards MJ, Richardson DJ, Paquete CM, Clarke TA. 2020. Role of multiheme cytochromes involved in extracellular anaerobic respiration in bacteria. Protein Science 29:830–842.

3. Lovley DR, Holmes DE, Nevin KP. 2004. Dissimilatory Fe(III) and Mn(IV) reduction. Advances in Microbial Physiology 49:219–286.

4. Salgueiro CA, Morgado L, Silva MA, Ferreira MR, Fernandes TM, Portela PC. 2022. From iron to bacterial electroconductive filaments: Exploring cytochrome diversity using Geobacter bacteria. Coordination Chemistry Reviews 452:214284.

5. Mishra S, Pirbadian S, Mondal AK, El-Naggar MY, Naaman R. 2019. Spin-dependent electron transport through bacterial cell surface multiheme electron conduits. Journal of the American Chemical Society 141:19198–19202.

6. Futera Z, Ide I, Kayser B, Garg K, Jiang X, van Wonderen JH, Butt JN, Ishii H, Pecht I, Sheves M, Cahen D, Blumberger J. 2020. Coherent electron transport across a 3 nm bioelectronic junction made of multi-heme proteins. J Physical Chemistry Letters 11:9766–9774.

7. Fu T, Liu X, Gao H, Ward JE, Liu X, Yin B, Wang Z, Zhuo Y, Walker DJF, Joshua Yang J, Chen J, Lovley DR, Yao J. 2020. Bioinspired bio-voltage memristors. Nature Communications 11:1861.

8. Terrell JL, Tschirhart T, Jahnke JP, Stephens K, Liu Y, Dong H, Hurley MM, Pozo M, McKay R, Tsao CY, Wu H-C, Vora G, Payne GF, Stratis-Cullum DN, Bentley WE. 2021. Bioelectronic control of a microbial community using surface-assembled electrogenetic cells to route signals. Nature Nanotechnology 16:688–697.

9. Lovley DR, Ueki T, Zhang T, Malvankar NS, Shrestha PM, Flanagan KA, Aklujkar M, Butler JE, Giloteaux L, Rotaru A-E, Holmes DE, Franks AE, Orellana R, Risso C, Nevin KP. 2011. Geobacter: The microbe electric’s physiology, ecology, and practical applications, p. 1–100. In Poole, RK (ed.), Advances in Microbial Physiology. Academic Press.

10. Levar CE, Chan CH, Mehta-Kolte MG, Bond DR. 2014. An inner membrane cytochrome required only for reduction of high redox potential extracellular electron acceptors. mBio 5:e02034–14.

11. Levar CE, Hoffman CL, Dunshee AJ, Toner BM, Bond DR. 2017. Redox potential as a master variable controlling pathways of metal reduction by Geobacter sulfurreducens. ISME Journal 11:741–752.

12. Zacharoff L, Chan CH, Bond DR. 2016. Reduction of low potential electron acceptors requires the CbcL inner membrane cytochrome of Geobacter sulfurreducens. Bioelectrochemistry 107:7–13.

13. Joshi K, Chan CH, Bond DR. 2021. Geobacter sulfurreducens inner membrane cytochrome CbcBA controls electron transfer and growth yield near the energetic limit of respiration. Molecular Microbiology 116:1124–1139.

14. Jiménez Otero F, Chan CH, Bond DR. 2018. Identification of different putative outer membrane electron conduits necessary for Fe(III) citrate, Fe(III) oxide, Mn(IV) oxide, or electrode reduction by Geobacter sulfurreducens. Journal of Bacteriology 200:e00347–18.

15. Wang F, Gu Y, O’Brien JP, Yi SM, Yalcin SE, Srikanth V, Shen C, Vu D, Ing NL, Hochbaum AI, Egelman EH, Malvankar NS. 2019. Structure of microbial nanowires reveals stacked hemes that transport electrons over micrometers. Cell 177:361–369.e10.

16. Aklujkar M, Coppi MV, Leang C, Kim BC, Chavan MA, Perpetua LA, Giloteaux L, Liu A, Holmes DEY 2013. 2013. Proteins involved in electron transfer to Fe(III) and Mn(IV) oxides by Geobacter sulfurreducens and Geobacter uraniireducens. Microbiology 159:515–535.

17. Wang F, Mustafa K, Suciu V, Joshi K, Chan CH, Choi S, Su Z, Si D, Hochbaum AI, Egelman EH, Bond DR. 2022. Cryo-EM structure of an extracellular Geobacter OmcE cytochrome filament reveals tetrahaem packing. Nature Microbiology 1–10.

18. Orellana R, Leavitt JJ, Comolli LR, Csencsits R, Janot N, Flanagan KA, Gray AS, Leang C, Izallalen M, Mester T, Lovley DR. 2013. U(VI) reduction by diverse outer surface c-type cytochromes of Geobacter sulfurreducens. Applied and Environmental Microbiology 79:6369–6374.

19. Gray HB, Winkler JR. 1996. Electron transfer in proteins. Annual Reviews in Biochemistry 65:537–561.

20. Volkov AN, Nuland NAJ van. 2012. Electron transfer interactome of cytochrome c. PLOS Computational Biology 8:e1002807.

21. de la Lande A, Babcock NS, Řezáč J, Sanders BC, Salahub DR. 2010. Surface residues dynamically organize water bridges to enhance electron transfer between proteins. Proceedings of the National Academy of Sciences 107:11799–11804.

22. Meschi F, Wiertz F, Klauss L, Blok A, Ludwig B, Merli A, Heering HA, Rossi GL, Ubbink M. 2011. Efficient electron transfer in a protein network lacking specific interactions. Journal of the American Chemical Society 133:16861–16867.

23. Fernandes AP, Nunes TC, Paquete CM, Salgueiro CA. 2017. Interaction studies between periplasmic cytochromes provide insights into extracellular electron transfer pathways of Geobacter sulfurreducens. Biochemical Journal 474:797–808.

24. Volkov AN. 2015. Structure and function of transient encounters of redox proteins. Accelerated Chemical Research. 48:3036–3043.

25. Sturm G, Richter K, Doetsch A, Heide H, Louro RO, Gescher J. 2015. A dynamic periplasmic electron transfer network enables respiratory flexibility beyond a thermodynamic regulatory regime. ISME Journal 9:1802–1811.

26. Lloyd JR, Leang C, Hodges Myerson AL, Coppi MV, Cuifo S, Methe B, Sandler SJ, Lovley DR. 2003. Biochemical and genetic characterization of PpcA, a periplasmic c-type cytochrome in Geobacter sulfurreducens. Biochemical Journal 369:153–161.

27. Ueki T, DiDonato LN, Lovley DR. 2017. Toward establishing minimum requirements for extracellular electron transfer in Geobacter sulfurreducens. FEMS Microbiology Letters 364:093.

28. Dantas JM, Brausemann A, Einsle O, Salgueiro CA. 2017. NMR studies of the interaction between inner membrane-associated and periplasmic cytochromes from Geobacter sulfurreducens. FEBS Letters 591:1657–1666.

29. Antunes JMA, Silva MA, Salgueiro CA, Morgado L. 2022. Electron flow from the inner membrane towards the cell exterior in Geobacter sulfurreducens: Biochemical characterization of cytochrome CbcL. Frontiers in Microbiology 13.

30. Pokkuluri PR, Londer YY, Yang X, Duke NEC, Erickson J, Orshonsky V, Johnson G, Schiffer M. 2010. Structural characterization of a family of cytochromes c7 involved in Fe(III) respiration by Geobacter sulfurreducens. Biochimica et Biophysica Acta (BBA) - Bioenergetics 1797:222–232.

31. Ding Y-HR, Hixson KK, Aklujkar MA, Lipton MS, Smith RD, Lovley DR, Mester T. 2008. Proteome of Geobacter sulfurreducens grown with Fe(III) oxide or Fe(III) citrate as the electron acceptor. Biochimica et Biophysica Acta (BBA) - Proteins and Proteomics 1784:1935–1941.

32. Shelobolina ES, Coppi MV, Korenevsky AA, DiDonato LN, Sullivan SA, Konishi H, Xu H, Leang C, Butler JE, Kim B-C, Lovley DR. 2007. Importance of c-type cytochromes for U(VI) reduction by Geobacter sulfurreducens. BMC Microbiology 7:16.

33. Santos TC, Silva MA, Morgado L, Dantas JM, Salgueiro CA. 2015. Diving into the redox properties of Geobacter sulfurreducens cytochromes: a model for extracellular electron transfer. Dalton Transactions 44:9335–9344.

34. Morgado L, Bruix M, Londer YY, Pokkuluri PR, Schiffer M, Salgueiro CA. 2007. Redox-linked conformational changes of a multiheme cytochrome from Geobacter sulfurreducens. Biochemical and Biophysical Research Communications 360:194–198.

35. Morgado L, Bruix M, Pessanha M, Londer YY, Salgueiro CA. 2010. Thermodynamic characterization of a triheme cytochrome family from Geobacter sulfurreducens reveals mechanistic and functional diversity. Biophysical Journal 99:293–301.

36. Morgado L, Dantas JM, Bruix M, Londer YY, Salgueiro CA. 2012. Fine tuning of redox networks on multiheme cytochromes from Geobacter sulfurreducens drives physiological electron/proton energy transduction. Bioinorganic Chemistry and Applications 2012:e298739.

37. Zacharoff LA, Morrone DJ, Bond DR. 2017. Geobacter sulfurreducens extracellular multiheme cytochrome PgcA facilitates respiration to Fe(III) oxides but not electrodes. Frontiers in Microbiology 8:2481.

38. Chan CH, Levar CE, Zacharoff L, Badalamenti JP, Bond DR. 2015. Scarless genome editing and stable inducible expression vectors for Geobacter sulfurreducens. Applied and Environmental Microbiology. 81(20):7178.

39. Cornell R, Schwertmann U. 2006. The Iron Oxides: structure, properties, reactions, occurrences and uses. Wiley-VCH Verlag GmbH & Co

40. Hallberg ZF, Chan CH, Wright TA, Kranzusch PJ, Doxzen KW, Park JJ, Bond DR, Hammond MC. 2019. Structure and mechanism of a Hypr GGDEF enzyme that activates cGAMP signaling to control extracellular metal respiration. eLife 8:e43959.

41. Stookey LL. 1970. Ferrozine---a new spectrophotometric reagent for iron. Analytical Chemistry 42:779–781.

42. Ross DE, Ruebush SS, Brantley SL, Hartshorne RS, Clarke TA, Richardson DJ, Tien M. 2007. Characterization of protein-protein interactions involved in iron reduction by Shewanella oneidensis MR-1. Applied and Environmental Microbiology 73:5797–5808.

43. Haider SR, Reid HJ, Sharp BL. 2012. Tricine-SDS-PAGE, p. 81–91. In Kurien, BT, Scofield, RH (eds.), Protein Electrophoresis: Methods and Protocols. Humana Press, Totowa, NJ.

44. Choi K-H, Mima T, Casart Y, Rholl D, Kumar A, Beacham IR, Schweizer HP. 2008. Genetic tools for select-agent-compliant manipulation of Burkholderia pseudomallei. Applied and Environmental Microbiology 74:1064–1075.

45. Zückert WR. 2014. Secretion of bacterial lipoproteins: Through the cytoplasmic membrane, the periplasm and beyond. Biochimica et Biophysica Acta (BBA) - Molecular Cell Research 1843:1509–1516.

46. Grabowicz M, Silhavy TJ. 2017. Redefining the essential trafficking pathway for outer membrane lipoproteins. Proceedings of the National Academy of Sciences 114:4769–4774.

47. Chan CH, Levar CE, Jiménez-Otero F, Bond DR. 2017. Genome scale mutational analysis of Geobacter sulfurreducens reveals distinct molecular mechanisms for respiration and sensing of poised electrodes versus Fe(III) oxides. Journal of Bacteriology 199:e00340–17.

48. Ding Y-HR, Hixson KK, Giometti CS, Stanley A, Esteve-Núñez A, Khare T, Tollaksen SL, Zhu W, Adkins JN, Lipton MS, Smith RD, Mester T, Lovley DR. 2006. The proteome of dissimilatory metal-reducing microorganism Geobacter sulfurreducens under various growth conditions. Biochimica et Biophysica Acta (BBA) - Proteins and Proteomics 1764:1198–1206.

49. Estevez-Canales M, Kuzume A, Borjas Z, Füeg M, Lovley D, Wandlowski T, Esteve-Núñez A. 2015. A severe reduction in the cytochrome c content of Geobacter sulfurreducens eliminates its capacity for extracellular electron transfer. Environmental Microbiology Reports 7:219–226.

50. Zhang X, Prévoteau A, Louro RO, Paquete CM, Rabaey K. 2018. Periodic polarization of electroactive biofilms increases current density and charge carriers concentration while modifying biofilm structure. Biosensors and Bioelectronics 121:183–191.

51. Esteve-Núñez A, Sosnik J, Visconti P, Lovley DR. 2008. Fluorescent properties of c-type cytochromes reveal their potential role as an extracytoplasmic electron sink in Geobacter sulfurreducens. Environmental Microbiology 10:497–505.

